# Ecological correlates of reproductive status in a guild of Afrotropical understory trees

**DOI:** 10.1101/2021.01.14.426416

**Authors:** Andrea P. Drager, Michael Weylandt, George Chuyong, David Kenfack, Duncan W. Thomas, Amy E. Dunham

## Abstract

The relative abundance patterns of tropical trees have been of interest since the expeditions of Alfred Russel Wallace, but little is known about how differences in relative abundance relate to reproductive patterns. Flowering is resource-dependent and fitness differences as well as differences in the quality of the abiotic and biotic neighborhood may contribute to the variation in reproductive status responsible for population-level flowering patterns. This variation determines the density and distance between flowering conspecifics and may alter relative abundance extremes among species during reproduction, factors known to influence pollination success. We collected flowering status data for a guild of twenty-three co-occurring tree species that flower in the understory of the Korup Forest Dynamics Plot in Cameroon. We examined how the occurrence and location of reproductive events were related to spatial patterns of adult abundance, focal tree size, neighborhood crowding, and habitat, while accounting for the influence of shared ancestry. Across species, the probability of flowering was higher for individuals of rarer species and for larger individuals but was unrelated to neighborhood crowding or habitat differences. Relative abundance extremes were reduced when only flowering individuals were considered, leading to a negative relationship between plot abundance and flowering probability at the species level that was not structured by shared ancestry. Spatially, flowering conspecifics tended to be overdispersed relative to all adult conspecifics. Rare species are predicted to suffer Allee effects or reduced fitness due to the difficulty of finding mates at low densities and frequencies. Here, however, rare species appear to maximize the size of their mate pool, compared to abundant species. If this partial ‘leveling of the playing field’ during reproduction is typical, it has consequences for our understanding of biodiversity maintenance and species coexistence in tropical forests.

## INTRODUCTION

The distribution of individuals in space and time is critical for a species’ reproductive biology, its use of resources, and how individuals interact with other species (Chesson, 2000; MacArthur, 1958; Wolkovich et al., 2014). In sedentary organisms, reproduction is also a unique opportunity to act beyond one’s immediate spatial neighborhood, via gene dispersal. Individuals of iteroparous, long-lived species such as trees may skip reproductive opportunities occasionally or frequently if certain resource-related conditions are not met (Bull & Shine, 1979; Isagi et al., 1997; Obeso, 2002; Waples & Antao, 2014). Whether and when individuals flower has direct consequences for reproductive success, demography, and genetic exchange (Ghazoul, 2005; Hendry & Day, 2005; Kang et al., 2003; Robledo‐Arnuncio & Austerlitz, 2006; Stacy et al., 1996) as well as for plant-pollinator interactions (Levin & Anderson, 1970; Kenta et al. 2004; Bell et al., 2005; Dietzsch et al., 2011), potentially influencing species coexistence and the persistence of rare species (Hart et al., 2016; Osada et al., 2003). High variation in flowering has been found in tropical and temperate tree communities, both among species and within populations, but few studies explore these patterns at the individual level in the context of reproductive strategies (Wright et al., 2005; Queenborough et al. 2007; Clark et al. 2010; Visser et al., 2016).

Species rarity or abundance in a community may alter the costs and benefits of temporal and spatial dispersion, particularly during events requiring phenological synchrony between individuals, such as reproduction (Elzinga et al., 2007; Velazquez-Castro & Eichhorn, 2017). In trees, flowering patterns in space and time are determined not only by the distribution and abundance of adults but by variation in the reproductive status among individuals. Individual flowering status may not scale up to species-level patterns in a community, either because there is temporal asynchrony among individuals during a reproductive cycle, or because there is variability in whether individuals participate in a cycle at all (Newstrom et al., 1994; Sakai et al., 1999; Boyle & Bronstein, 2012; Bush et al., 2016;). Patterns of reproductive heterogeneity within a population may be especially important for species that require outcrossing for reproduction, such as many woody species, since the number of flowering individuals represents the maximum potential number of effective breeders (Bawa et al., 1985; Murawski et al., 1991).

Plant reproduction is generally limited by two factors: the amount and the quality of pollen received, and the energetic resources available for reproduction (Ashman et al., 2004; Rosenheim et al., 2014). Rarer species may face greater pollination challenges, potentially leading to Allee effects, defined as reduced fitness due to the difficulty of finding mates at low densities (Asmussen, 1979; Kunin, 1997; Vamosi et al., 2013). For co-flowering species that share pollinators, the effects of low density may be compounded by being at a low flowering frequency in the community, leading to reproductive interference from more abundant species (Whitney, 2009). Resource distribution is a strong predictor of pollinator visitation (Kunin & Iwasa, 1996; Schmid et al., 2015) and the clustered flowering of individuals may improve the pollination outcomes for rarer species. Individuals of abundant species may instead reduce the fitness costs of intraspecific competition for pollinators by flowering asynchronously or by having a percent that abstains altogether (Momose, 2004). Flowering patterns have implications for offspring fitness as well, though often in species-specific ways. Pollen flow is highest among near-neighbors but the strong spatial genetic structure seen in many tree populations suggests that reproduction between near neighbors may have negative fitness consequences for offspring (Hardy et al., 2006; Jones & Comita, 2008). At the other extreme, isolated trees, while frequently still connected, may receive less outcross pollen from fewer pollen donors (Murawski et al., 1991; Dick et al., 2008).

A more proximate driver of flowering patterns may be a tree’s response to resource availability, as mediated by local abiotic and biotic factors and climatic influences. Climate-related drivers (eg. rainfall, solar radiation, temperature, CO2) are triggers of the onset of flowering and are associated with interannual covariance in community-level flowering patterns (Schauber et al.2002; Pau et al. 2017). However, within communities, variation exists in the magnitude, or the proportion of a population that flowers in a given year (Thomas & LaFrankie, 1993; Sakai et al., 1999; Queenborough et al., 2007). Flowering is resource-dependent, and whether an individual tree has the resources to participate in a reproductive bout may depend on the amount of resource capture and allocation in the previous season as well as the species-specific timing of allocation to floral development (Hoch et al., 2013; Sedgley & Griffin, 2013; Blake-Mahmud & Struwe, 2019). Spatial differences in nutrients and moisture will also mediate individual resource availability for species found across gradients in habitat quality (Libalah et al. 2017). Additionally, larger individuals tend to have more access and ability to store resources, and therefore may be less likely to skip reproductive opportunities, though this may not be the case for all species (Wright et al., 2005; Queenborough et al., 2007; Dainou et al., 2012; Visser at al.,2016; Bruijning et al., 2017). However, at the largest size classes representing the oldest trees, senescence and increased pathogen burden may reduce reproduction (Clark et al., 2010; Mothe et al., 2017). Some species are able to allocate some resources to flowering yearly, even in resource-poor years when there is heavy abortion of immature fruit; other species show intermittent, often irregular flowering (Obeso, 2002).

Individual resource availability can be influenced by competition with neighbors for access to limiting resources such as light, water, and soil nutrients. Much previous work has quantified the effect of neighbor competition on growth and survival of trees using a spatially-explicit approach that incorporates the size, distance and in some instances, identity of neighbors around a focal tree (Bella, 1971; Canham et al., 2004; Lasky et al. 2015; Taylor et al., 2017). While theory assumes stronger competitive effects of increased density among conspecifics than among heterospecifics, due to more similar resource requirements, this pattern is less evident in tropical forests, where the majority of species are low-density and nearest non-seedling neighbors are more likely to be heterospecifics (Uriarte et al., 2004). A few studies have extended this neighborhood competition approach to reproduction, specifically fruiting, with results suggesting that trees experiencing higher crowding may be less likely to fruit (Minor & Kobe, 2019) or have lower reproductive output than those that are less crowded (Haymes & Fox, 2012; Nottebrock et al., 2017). However, as seen in orchard and agroforestry systems, reduced reproductive output can also lead to more regular flowering, a result of avoiding the resource sink of heavy fruiting that leads to intermittent breeding in some species (Sedgley & Griffin, 2013; Vaast et al., 2006). Resource limitation, as mediated by abiotic or biotic factors may impact flowering status in ways that vary somewhat both among and within species.

Tropical tree communities are interesting systems for examining individual variation in flowering status across species. These communities are highly biodiverse, with species abundances that vary by several orders of magnitude; in addition, the vast majority of tropical tree species rely on animal mutualists to carry their pollen between conspecifics (Bawa, 1990). In this study, we used flowering data collected during a peak flowering season in the 50 ha Korup Forest Dynamics Plot in Cameroon to explore how the occurrence and location of individual flowering events relate to adult abundance patterns. Then we asked whether we could identify a signature of resource-related drivers of these flowering patterns: we modeled the flowering status of individual trees as a function of local species abundance, tree size, neighborhood crowding, and fine-scale habitat, while assessing how much variation among species was due to shared ancestry or simply to species differences unaccounted for in our models. We predicted that, within species, if flowering is sensitive to endogenous resource availability and competition associated with crowding, both larger and less crowded individuals will be more likely to flower. We also expected that flowering probability would be mediated by habitat differences among individuals, particularly if species are relatively habitat-specialist. At the species level, we expected that if rarer species benefit from aggregated flowering, we would see this both spatially, in terms of clumped distributions of flowering individuals, and in terms of the proportion of trees that flower.

## METHODS

### Site

Over the past decades, the establishment of long-term forest dynamics plots large enough to include rare species has greatly increased our ability to obtain individual-level data in highly biodiverse communities. One such plot is the 50 ha Korup Forest Dynamics Plot (KFDP) in Korup National Park, Cameroon. Established in 1996, the KFDP is part of a collaborative global network of long-term forest monitoring plots affiliated with the Smithsonian Forest Global Earth Observatory (ForestGEO). All stems >1cm in diameter at breast height (DBH) have been mapped, measured and identified (Thomas et al., 2003). This lowland tropical mixed forest is part of the Guineo-Congolian forest zone; the KFDP itself contains ~494 woody species (Thomas et al., 2003). Patterns of relative abundance are similar to those of other diverse tropical forests (Kenfack et al., 2007).

Strong seasonality at this site produces a distinct flowering period triggered by the end of the dry season, making it ideal for studying flowering in a biodiverse community. Most species flower between the months of February and April, as rainfall becomes increasingly heavy (D.W. Thomas, pers.comm.). The plot is a mature closed-canopy forest with disturbance limited to rare gaps produced by wind-falls (Egbe et al., 2012). It is divided by a creek running east to west that roughly delimits the steeper topography in the northern half from that of the flatter southern half (Chuyong et al., 2011). These topographical differences between the north and south of the plot also correspond to a difference in species distribution patterns for many species (Kenfack et al., 2014). Due to logistical constraints, data was collected in the 25 ha south of the creek where habitat is classified according to Chuyong et al. (2011) into two types based on slope, elevation, and convexity: “low flat” and “low depressions”, and the “low depressions” habitat is inundated during the wet season (Chuyong et al. 2011). All focal species had individuals present in both habitat types, and all data presented in this study are from this southern 25 ha only.

### Field Methods

During a period of five weeks from February 29th-April 5th, 2016, we recorded the tree tags of all reproductive individuals that flowered within approximately four meters of the ground. This distance was the cut-off for how far we could clearly observe buds and flowers on the tree without the use of binoculars. The species in our dataset are therefore composed of a guild of share-tolerant trees that produce flowers at the base of or along the trunk, on low branches, or that are shrubby; the tallest species are <15m and are cauliflorous. In order to thoroughly sample the reproductive phenology of those individuals active at the tail end of our sampling period, we recorded budding as well as flowering. The time from when buds are first visible to the completion of flowering is a process of at least several weeks. By including budding phenology, we are confident we have captured those species and individuals that flower towards the end of the season. Therefore, the reproductive phenology data analyzed here are grouped into one binary variable denoting reproductive status; it represents co-flowering within the 2016 flowering season, but not necessarily direct overlap at the level of individual trees. We exhaustively searched the 25ha study area south of the stream that runs roughly West to East, working our way East to West and North to South through the demarcated 1ha subplots, sampling approximately two subplots a day, depending on how easily we could move through the vegetation (patches with dense, swampy terrain and spiny lianas took longer). We were able to thoroughly census each subplot at least twice during the five-week period, and by including both budding and flowering stages in our reproductive status metric, we were able to document at least one of these stages even for those species with the shortest flowering phenology (eg. males of *Crotogynopsis sp.nov.*).

### Species Selection

There were 169 woody species with at least one individual over 1cm DBH (and thus included in the KFDP data) that are either understory species or that flower in the understory (e.g. cauliflorous). Of these, about half (n=90) had at least one reproductive individual in 2016. Focal species are summarized in Table S1, a summary of the full reproductive census data is in Table S3, and visual summary is in Fig. S2. From our dataset of individual observations, we removed species with cryptic flowers, for fear they had been under-sampled (such as *Uvariopsis korupensis*), thus the dataset is biased towards more conspicuous (easy to see or strongly scented) species. We also removed species that had some shorter individuals flowering within the 4m cut-off, but where the majority tended to flower higher up in the canopy (such as *Cola rostrata*). Species also removed are the smaller shrub species (such as *Phyllobotryon spathulatum*), where we encountered >10% of flowering individuals untagged; this indicated that adults are not well-represented in the >1cm DBH KFDP data and would, therefore, incorrectly classify such species as much less abundant than they actually are. Lastly, for statistical reasons, we removed those species with fewer than three individual records of budding or flowering during the five-week study period (n=46 species; such as *Trichoscypha acuminata*). This cautious culling of uncertain species records as well as those with too few flowering individuals resulted in a sample size of 1782 records of 23 species.

After identifying our focal species, we created an additional dataset of total adults for each species in the 25ha, using the most recent census dataset available (2008-2009) from KFDP (Fig. 1). The definition of an adult for each species was species-specific and was defined as individuals having a DBH greater or equal to the minimum diameter of an individual in the observed flowering cohort of each species in 2016. This is a conservative estimate, as the second census was gathered in 2008-2009, representing a seven-year growth interval. Our data does not account for any demographic changes in abundance (death or recruitment of adults) that may have occurred in this interval. New reproductive recruits, which were not in the 2008-2009 census, were not included in the analysis because we do not have comparable data on non-reproductive individuals of those size classes. Otherwise, any bias introduced is likely in the direction of misidentifying dead adults as live, non-flowering individuals.

**Figure 1.**
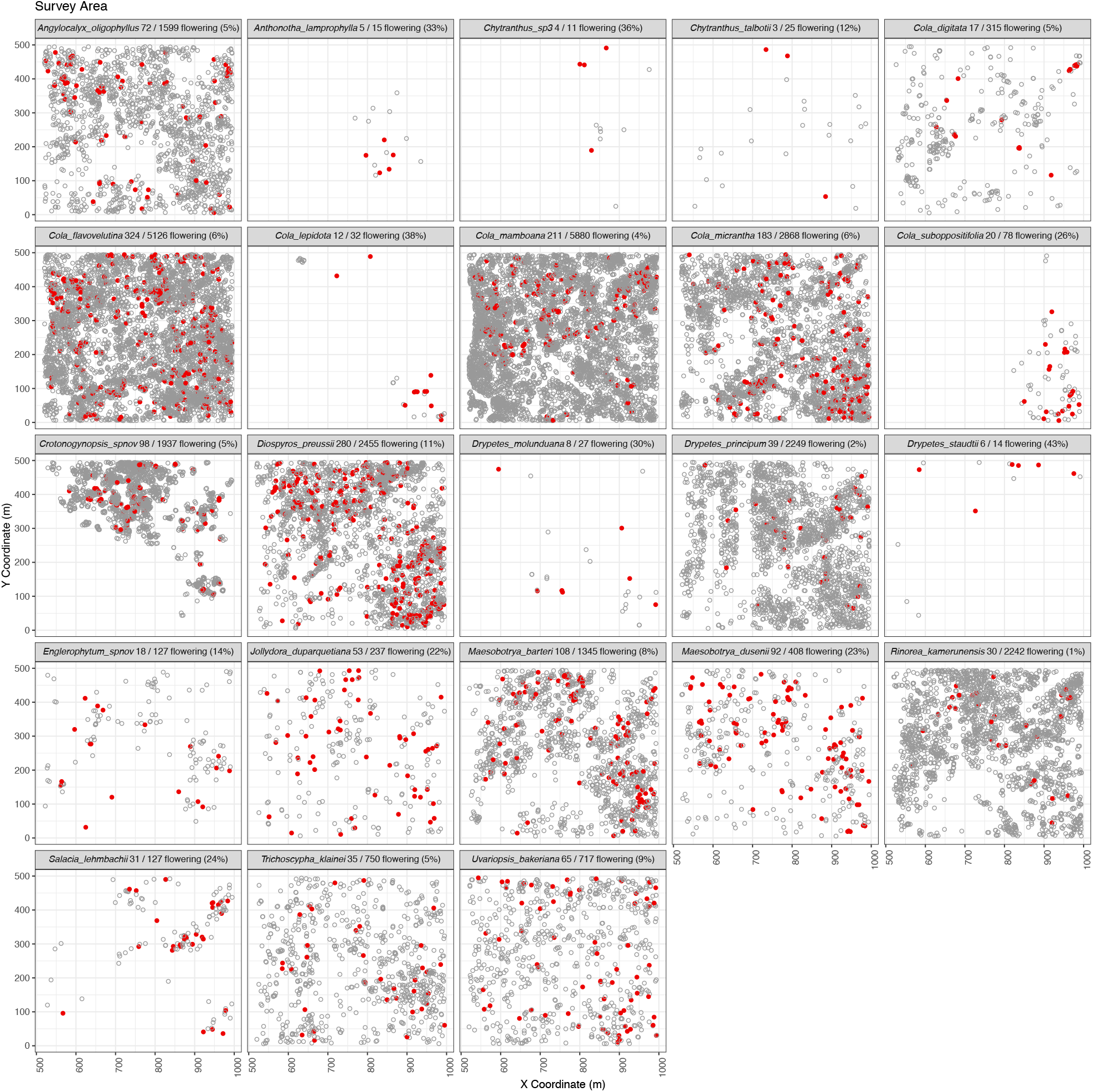
Species maps of flowering (red) and non-flowering (gray) adults in the 25 ha survey area. Fractions and percentages flowering are also recorded.

### Analysis

All statistical analyses were done in R version 3.4.3 (R Project Core Development Team, Vienna, Austria). To fit the models described in the next section, we used the `brms` package (Bürkner, 2017), which fits a wide range of generalized linear mixed models in a Bayesian framework. The `brms` package is built upon the Stan probabilistic programming language (Carpenter et al., 2017), which allows for efficient sampling using a variant of the No-U-Turn Sampler scheme for Hamiltonian Monte Carlo (Neal, 2011; Hoffman & Gelman, 2014; Betancourt, 2017). Additionally, the package `spatstat` (Baddeley, Rubak, & Turner, 2015) was used to calculate nearest-conspecific distances and neighborhood crowding indices, the package pROC (Robin et al., 2011) was used to calculate the AUC metric used for model comparison and the `bayesplot` (Gabry & Tristan Mahr, 2018) and `ggplot2` (Wickham, 2016) packages were used to visualize our results.

### Ecological correlates of flowering status

To investigate the effects of local abundance, tree size, neighborhood crowding and habitat at different spatial scales on flowering status, we fit a set of generalized linear mixed models in a Bayesian framework using the `brms` package V.2.8.0 (Betancourt, 2017; *brms*, n.d., 2017; Carpenter et al., 2017; Hoffman & Gelman, 2014; Neal, 2011). Local abundance, a species-level term, is the abundance of adults of a species in the sample area. Tree size is the diameter at breast height of a tree, obtained from the 2008-2009 KFDP census. The neighborhood competition index (NCI), or crowding index, consists of the squared diameter (DBH) of each neighbor tree divided by the square of its distance to the focal individual, summed over all neighbors (Lasky et al 2015; Taylor et al 2017). We calculated NCI for three concentric radii around each focal individual: 5m, 10m, 15m, using a buffer of equal size around the entire study area to ensure the completeness of neighborhoods for individuals located near the plot edge. We selected these spatial extents based on previous work on competitive effects between adults in tropical forests (Lasky et al., 2015) as well as data on crown diameters for rainforest species (Osunkoya et al., 2007). We hypothesized that the use of nested spatial extents could allow us to observe at what distance, if any, neighbors had the strongest impact.

Due to the phylogenetic diversity present in our species sample, we expected variation among species in the relationships between our covariates and flowering status, as well as the potential for unexplained variation among species. To address species-specific responses among covariates, we fit species random slopes for each individual-level covariate. In order to understand whether variation in flowering status among species was structured by shared ancestry, we compared models with and without a parametric phylogenetic component. If certain clades had higher proportions of flowering adults than others, then we expected to find stronger support for the model with variance structured by phylogenetic distance between species (Harvey & Pagel, 1991). Similarly, if phylogenetic structure was important to explaining the effect of certain covariates, we expected to see this manifested in terms of increased model fit in the phylogenetic random slopes model. The dated molecular phylogeny for the KFDP species was constructed by Ingrid Parmentier and Olivier Hardy et al. and provided by them upon request (Parmentier et al., 2013); from this, we created a distance matrix using phytools (Revell, 2012).

Our dataset was large enough to assess each model’s out of sample performance, so we split the data using a sampling scheme stratified by species into a training set consisting of 70% of the data and a test set consisting of 30% of the data ten times (see analysis code in SI7). The out of sample area under the ROC curve (“AUC”) was calculated and averaged over the ten splits for each model. AUC provides a more coherent measure of the accuracy of a probabilistic classification algorithm than simple misclassification error, particularly on severely unbalanced datasets such as ours, by assessing the tradeoff between sensitivity and specificity on a global basis, rather than at any particular cut-off. Our model achieved an out-of-sample AUC of approximately 71%, which compares favorably to black-box methods such as random forests (Brieman, 2001), which achieve an AUC of approximately 65% on the same data (Fig. S3). Our model (below) is a hierarchical (“mixed-effects”) logistic regression model, incorporating species-specific intercepts and slopes for each of the covariates used (Gelman & Hill, 2007). We note that our model did not make direct use of the phylogenetic relationships among sampled species as including this information did not notably improve the predictive performance of the resulting model. We also note that the predictive performance and estimated effect sizes of our model were robust to the particular choice of covariates used and additional robustness checks are presented in the supplementary materials.

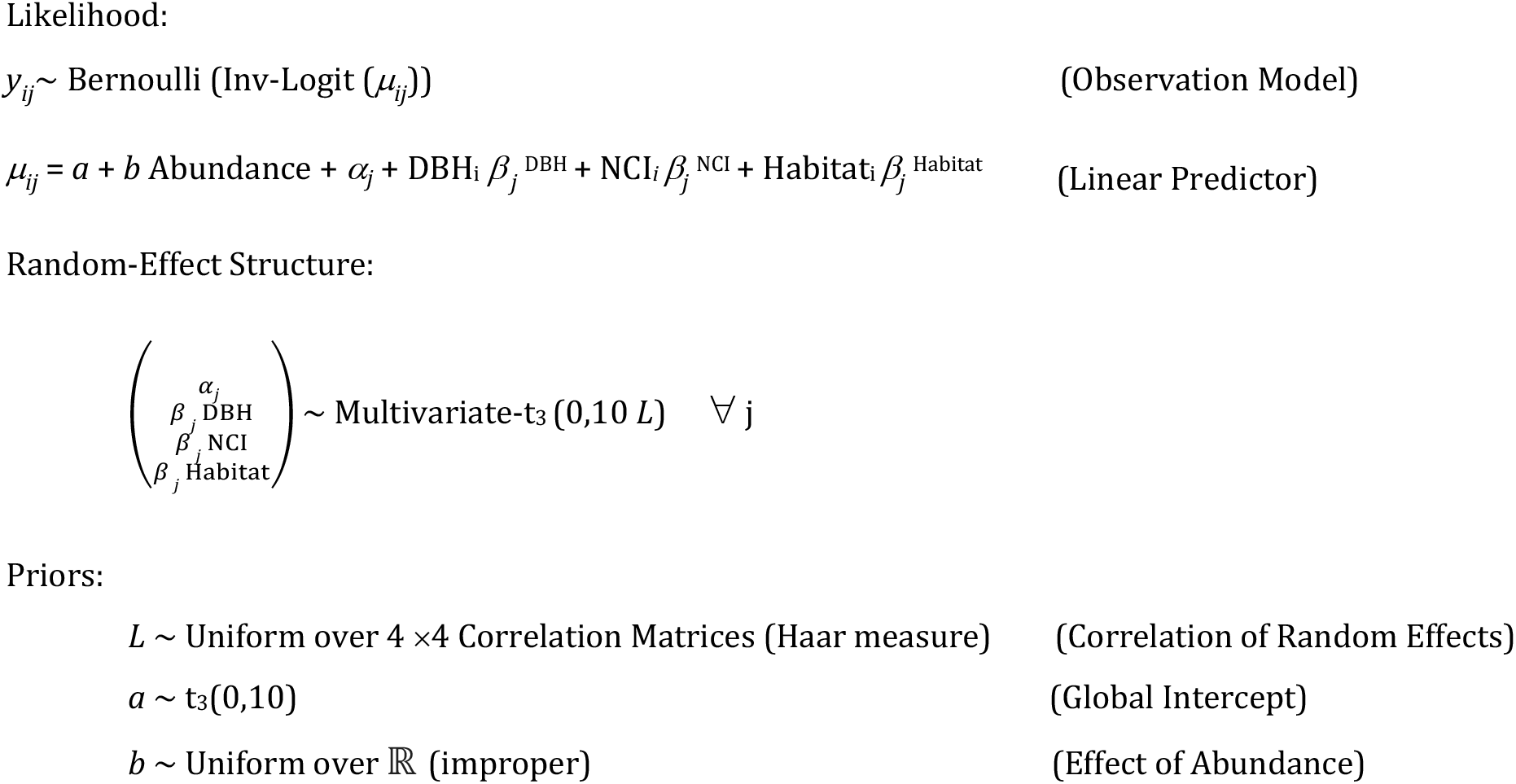

Where *i-*subscripts denote individual trees and *j*-subscripts denote tree species and where Mutivariate-t_3_(0,10,L) denotes a multivariate Student’s t-distribution with 3 degrees of freedom, mean 0, marginal standard deviation 10, and correlation L. (Note that in practice, a slightly different but numerically equivalent formulation is used internally by `brms` to improve smapling performance.)

Continuous predictor variables (other than the phylogenetic distance matrix) were z-transformed on a per-species basis to standardize them (Gelman et al., 2013). Our data exhibit minimal correlation between the (transformed) DBH, NCI and Habitat measurements, with a full-sample correlation of approximately −2.6%, allowing for straightforward interpretation of the marginal effects. Per-species correlations are reported in the supplementary materials (Fig. S5). For all parameters, the default priors of the `brms` package were used. The sampler was able to efficiently explore the posterior, and all MCMC convergence diagnostics were well within acceptable levels: the Gelman-Rubin-Brooks potential scale reduction factor (“R-hat”) statistic was between 1.000 and 1.004 for all parameters, the estimated effective sample size was over 1200 for all parameters of interest, and no divergent transitions were encountered (Gelman et al., 2013). Fixed parameters with 90% credible intervals that did not overlap zero were assumed to be significant.

### Spatial patterns of flowering

To understand whether the spatial pattern of flowering individuals was clustered, we compared nearest neighbor distances between flowering individuals and adults as a whole. We took 1000 random samples from the total adult population in the study area, for each species, with a sample size based on the number of flowering individuals. Trees were sampled randomly from the unique spatial distribution of each species using the *ranp* function in `spatstat`. Next, we calculated nearest neighbor distances for all individuals in both the observed flowering and the 1000 random samples. We chose nearest-neighbor distance from among the possible metrics of aggregation because it represents a measure that is likely relevant to pollinator foraging (Ghazoul, 2005; Kunin & Iwasa, 1996). We used a binomial sign test to assess the statistical significance of the differences in nearest neighbor distances of flowering individuals to the median distance for random samples of total adults of each species. We also used a one-sample t-test to test the pattern of clustering of flowering across species relative to the null expectation.

## Results

### Reproductive abundance different from adult abundance

In no case did all adult individuals of a species flower, and the proportion of adults of a species that were reproductive decreased with local abundance, irrespective of taxonomic family (Fig. 1; Fig. S1). The mean value for percent of adults of a species flowering was 16%, median 10%, range was 1.3%-50%.

### Abiotic and biotic predictors of flowering

At the community level, the strongest predictor of flowering was diameter at breast height (DBH), with the odds of an individual flowering increasing by approximately 22% with every cm gain in diameter, averaged across all species (Fig. 2). Adult abundance in the study area was also a non-trivial effect, with an increase in 1000 individuals representing a 36% decrease in the odds of flowering, all else being equal (Fig. S4). Within species, DBH was a significant positive predictor of flowering in 11/23 species, and a negative predictor for 2/23 species. There was no significant effect of neighborhood competition (NCI) on flowering among or within species at any of the three neighborhood spatial extents: 5m, 10m, 15m radii. We present the results of the 5m neighborhoods because this spatial extent is likely to be most relevant, given the small size of the trees in our study (Fig. 2). Habitat was also weakly associated with flowering status across species, with only 3/23 species exhibiting a significant negative effect associated with the low-depression habitat.

**Figure 2.**
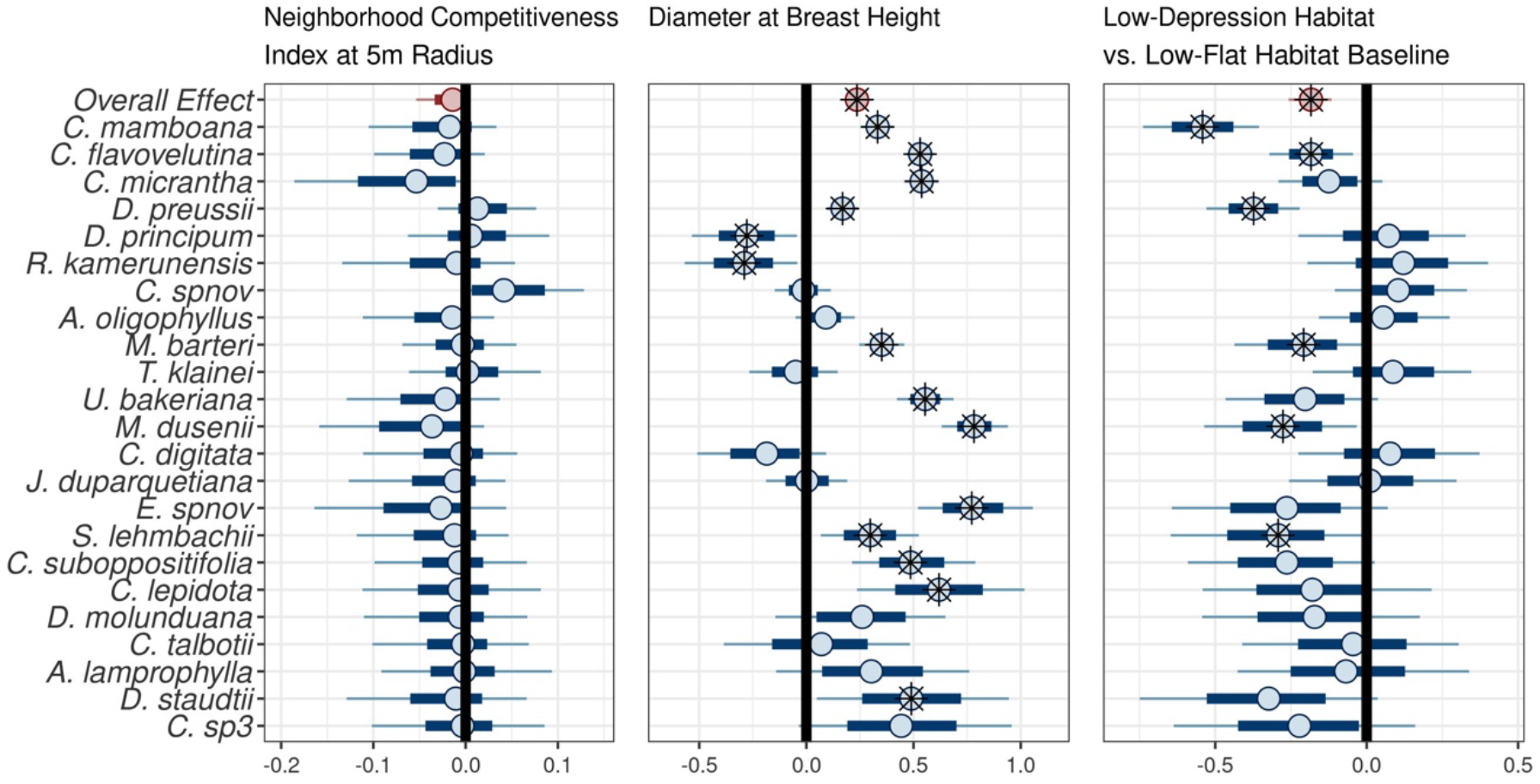
Median posterior probability estimates for species-level effects and weighted means of the per-species medians (Overall Effect, top), from panels left to right: NCI (5m radius neighborhoods), DBH, and Habitat on log-odds of flowering. The effect of DBH is clearly strongly positive for the majority of species sampled, while the effects of NCI and habitat are less pronounced. Coefficients correspond to Model 18 in Table S1. Results from 10m and 15m radii are nearly identical. Credible intervals that do not cross zero indicate significance at the 90% level and are starred. Species are ordered by adult abundance, from most abundant (top) to least (bottom).

Variation among species was a key feature of this data. Our best-performing model incorporates species-specific intercepts as well as species-specific responses to the DBH, NCI and Habitat covariates. Allowing for species-specific baseline flowering rates and species-specific sensitivities to covariates was critical to improving model performance above the predictive power of Random Forest (Table S2 & Fig. S3).

### Spatial patterns of flowering: overdispersion, not clustering

Across species, flowering individuals tended to be further from one another than adult trees were from one another. When flowering nearest-neighbor distances were compared to median distances for 1000 random samples of adults of each species, there was a significant overall pattern of over-dispersion among flowering individuals across species (t = 5.27, df = 22, *P* < 0.001) (Figure 3). This pattern of overdispersion was significant within species for fifteen of the species, predominantly the more abundant ones. Six of the species had flowering individuals at distances that were not different than those of adults in general, and in two cases, a wide spatial range in flowering distances coupled with low sample sizes produced over-dispersed but non-significant median values using the binomial sign test. Median flowering nearest-neighbor distances for all species ranged from 3.5m – 57m, and adult median nearest neighbor distances ranged from 1.5m – 49m, though one rarer species, *Cola lepidota*, was unusually clumped, with median distances between nearest neighbors similar to those of the most abundant species. Species in the Malvaceae (all in the genus *Cola*) clustered in terms of nearest neighbor distances despite the rarest and the most common having adult abundances differing by two orders of magnitude. Notably, no flowering patterns were significantly clumped relative to adult sample medians. Additionally, within-species correlations between tree size and distance to nearest conspecific neighbor were low and most were not significant (median correlation 0.00; range - 0.38, .033), and the direction of the relationship was variable, indicating that spatial flowering patterns of individuals do not simply represent the subset of spatial distributions of larger DBH individuals.

**Figure 3.**
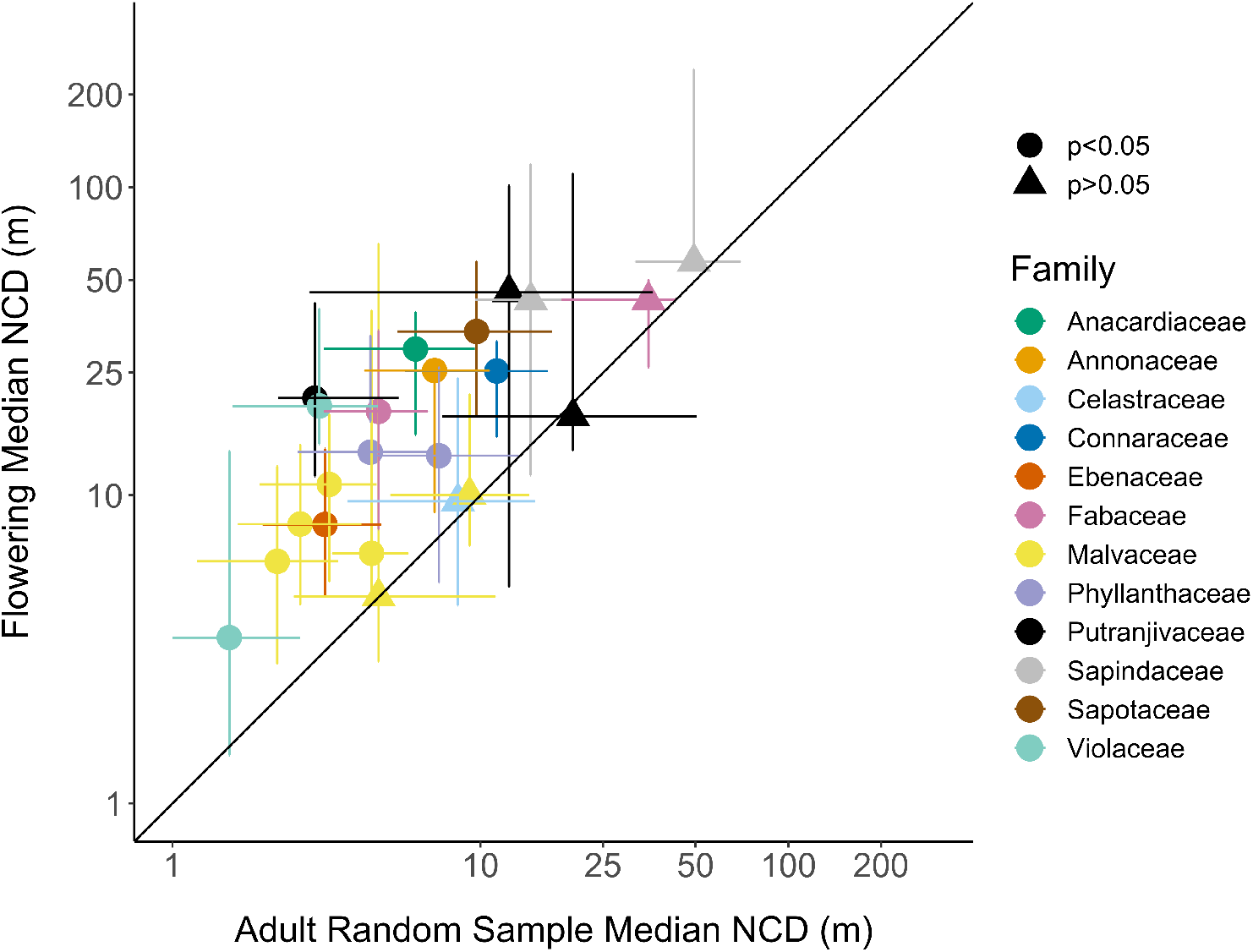
Median flowering nearest conspecific distances (NCD) plotted against median nearest conspecific distances from adult random samples, both with 25% and 75% quantiles, log transformed. Each point represents a species. The black line through the origin indicates values where nearest conspecific distances are equal for flowering individuals and for random conspecific adults. Species above the line have flowering individuals overdispersed relative to adult conspecifics.

## DISCUSSION

Variable, often low, participation in reproductive opportunities appears to be a feature of tropical trees that has the potential to influence both population and community dynamics in these biodiverse forests. However, these patterns have rarely been studied in an individually-based, spatially explicit, and comparative way. Here we described patterns of individual variation in reproductive status observed among a guild of trees that flower in the understory of a Guineo-Congolian rainforest, and we examined them from the perspective of reproductive ecology. We found that the probability of flowering was higher for rarer species and for larger individuals, and that flowering conspecifics were less spatially-aggregated than adults. For most species, flowering status was unrelated to neighborhood crowding or habitat type, proxies of local resource availability. Phylogenetic relationships did not explain the high variation in reproductive status among species; rather, there was a negative correlation between flowering and abundance even within taxonomic families. Our data show that intraspecific variation in flowering leads to a much narrower difference in flowering abundance among species, compared to patterns of adult relative abundance. This may result in a pollination landscape that is more favorable to the reproduction of low-density species than might be assumed from adult abundance patterns alone, with implications for coexistence dynamics and the persistence of rare species.

While no study we are aware of has examined the proportion of individuals flowering in relation to patterns of relative abundance in a community, a few studies have assessed the reproductive status for all adults in a given spatial extent in tropical forests: mean of 47% in small-canopy trees in deciduous moist forest, Panama (N=16; Wright et al. 2005); 33% in treelets-canopy trees in evergreen moist forest, Malaysia (N=21, year=1990; Thomas 1996), 26% in Myristicaceae in evergreen moist forest, Ecuador (N=16, year=2003; Queenbrough et al. 2007) compared to 16% in small trees and treelets in evergreen moist forest, Cameroon (N=23; this study). Community means are expected to fluctuate interannually due to changes in climate, so snapshot comparisons are not very informative, however, what these numbers do illustrate is the low overall participation of tropical trees in a given reproductive bout.

A decrease in the number of potential flowering individuals on the part of abundant species may be to the advantage of rarer species when competing for pollination services. If higher relative flowering abundance (frequency) in the community is associated with increased pollinator preferences or encounter rates, then a more even distribution of flowering individuals across species is expected to benefit the rarer species. For example, *Chytranthus sp.3*, the rarest species in our sample, with 12 adult individuals has a relative abundance of 12 out of 29,609 adult focal trees in the community (4.05 E-4). However, if you consider the relative abundance for that same species, based only on flowering individuals in our sample, the value is five and half times greater (four out of 1782 trees or 6.73 E-3). Smaller proportions of flowering individuals among the more abundant species could somewhat level the pollination playing field between rare and common species by lowering fitness costs for rarer species due to interspecific competition for pollinators. Additionally, a reduction in reproductive effort on the part of abundant species for whom mating opportunities abound could also be adaptive, increasing rather than reducing long-term individual reproductive output by allowing resource stocking (Bull & Shine, 1979), and reducing interspecific competition and inbreeding. In order to determine the significance of these reproductive patterns more broadly, in terms of coexistence and the persistence of rare species, we would need to both understand whether the trend we observed is consistent over multiple reproductive cycles, and how relative flowering abundance relates to fecundity in this community.

Flowering conspecifics tended to be less clustered relative to adult spatial patterns overall, and dioecious species were no exception, being present in our dataset at both lower and higher abundances. Clustered flowering has been suggested as adaptive, particularly for rare or dioecious species, based on the idea that proximity results in higher pollination success (Bawa et al., 1985; Ghazoul, 2005). However, whether differences of several meters vs. tens of meters affect pollination outcomes will depend on pollinator foraging behaviors as well as the ability of the tree to attract pollinators over given distances. While pollen flow in animal-pollinated trees appears to be highly context- and species-dependent, clustering can increase pollen receipt, though often with tradeoffs in terms of offspring fitness (Stacy et al., 1996; Dick et al., 2008; Jones and Comita, 2008). In small trees and herbs, pollen flow distances can range from tens to hundreds of meters, often traveling far beyond nearest neighbors (Dick et al., 2008; Dyer et al; 2012; Ottewell et al., 2012; Côrtes et al., 2013; Castilla et al., 2016; Grant et al., 2019); 50m is roughly the median distance separating flowering individuals of the lowest-density species in our study. If adequate pollination is occurring, a lack of nearest-neighbor flowering synchrony could be adaptive, reducing the bi-parental inbreeding that typically results from nearest-neighbor mating (Nason et al. 1998; Winn et al., 2011; Duminil et al., 2016).

Flowering is resource-dependent and larger trees have greater access to resources via larger crowns, higher canopy placement, and more extensive root systems. However, the relative size at the onset of maturity has been found to vary among tropical tree species, irrespective of life history parameters such as life form or shade tolerance (Ouedraogo et al. 2018; Thomas, 1996; Wright et al., 2005). Some species may trade-off growth for more frequent and earlier reproduction, suggesting that variation in this life history strategy among species may contribute to species coexistence (Thomas, 1996; Wright et al., 2005). As in previous studies, we found tree diameter (DBH) was the strongest predictor of flowering across species, though effect estimates were quite variable for some species and were not important for all of them (Figure 2). Low correlation among covariates allowed us to accurately estimate the effect of each predictor on flowering and to avoid the interpretational difficulties posed by strong collinearity. In particular, our results suggest that the effect of DBH was unrelated to other resource-related factors we studied, namely neighborhood crowding and habitat differences. It is possible that the strength of the flowering-DBH relationship estimated in any one year may be lessened by the phenomenon of “alternate bearing” or skipping reproductive opportunities that may follow a significant fruiting event the previous year. Variability in the flowering-DBH relationship may be associated with beginning reproduction at a smaller stature (Fig. S6), something feasible for our study species in terms of their reproductive ecology. Because they flower in the understory, many from their trunks and branches, these species are not constrained by the need to attain a certain height in order to reach pollinators in the canopy (Thomas 1996). Other traits involved in resource acquisition, such as specific leaf area, may also be useful to include as potential predictors of reproductive status is future studies (Visser et al., 2016).

In addition to tree size as an endogenous proxy for resource availability, we included proxies for the effect of biotic and abiotic influences. Estimating the effect of biotic interactions via neighborhood crowding (NCI), in terms of the density of neighboring trees and their sizes and distances to a focal tree, we found no correlation with flowering probability at any of the three spatial extents measured, either among or within species. NCI may not be a sensitive enough proxy in our study, because it was calculated based on the 2008-2009 KFDP census data, rather than on 2016 conditions (Detto et al. 2019). Light availability and its effect on carbohydrate stocks could be a key factor affecting reproductive patterns in understory trees and including direct measures of light availability may have improved our predictive power (Berdanier & Clark, 2016; Kainer et al., 2007; Wright et al., 2005). This neighborhood metric could also potentially be improved by incorporating information on the functional similarity of neighbors (Lasky et al., 2015).

Alternatively, it is possible that we saw no crowding effect on flowering status because flowering represents a less resource-intensive investment than fruiting, as manifested by the high rates of flower abortion seen even in cases where pollination was not limiting (Bawa & Webb, 1984; Armstrong & Irvine, 1989; Pearse et al. 2015). The majority of our focal species produce large-to medium-seeded, fleshy fruit that likely require a much greater resource investment than their flowers. Studies that have measured the impact of neighborhood interactions on reproduction in natural tree communities have focused on fruiting status and/or fruit crop size, rather than flowering status, and find both positive (Castilla et al., 2016; Jones & Comita, 2008) and negative effects of conspecific crowding on fecundity (Nottenbock et al., 2017; Haymes & Fox 2012), and crowding from all neighbors on fruiting status (Minor and Kobe, 2019). However, fruiting and fecundity are more complex because they depend on pollen receipt as well as on resource availability; the relative contributions of each of these factors to the crowding effect is unclear in several of these studies. Additionally, our study is limited to one reproductive season and therefore we were unable to distinguish between those individuals that were non-reproductive due to chronic resource limitation and those that were only temporarily non-flowering (“off-year”) following a significant fruiting event in the previous year. A longer, multi-year study would be able to more accurately assess the effects of crowding over multiple reproductive opportunities for these species; we leave this more detailed examination to future studies.

Habitat as a proxy for resource-related differences in the abiotic environment is conditional on how specialized a species is to certain abiotic conditions. Here habitat was based on topography, which is strongly correlated with moisture and nutrient levels in this part of the plot, with low-depressions being wetter and more nutrient poor than low-flat areas (Libalah et al. 2017). We found habitat was an important influence for three of the most abundant species, two of which are known to be narrow endemics *(Cola mamboana and Cola flavovelutina)*, though not specialists to either habitat (Kenfack et al., 2006; Chuyong et al., 2011), the third (*Diospyros preussii*) had more individuals flowering in the low-flat habitat but was not more abundant here compared to the low-depression habitat. For the remaining twenty, habitat was not strongly associated with flowering status. This lack of strong habitat preferences for flowering supports our decision to model species abundance at the level of the entire 25ha study area.

The lack of strong links between flowering status and crowding and habitat as proxies for local resource-related drivers suggests that future work may benefit from a more mechanistic approach. The success of resource-budget models in furthering our understanding of drivers in wind-pollinated, masting trees has created new synthetic directions for research (reviewed in: Pearse et al., 2016). Flowering dynamics in beech, for example, have been successfully predicted as the combined effects of climate cues and pollen and nutrient availability, although flowering intensity was not correlated with distance between conspecifics, and trees of similar sizes were not more likely to flower the same year (Abe et al., 2016). Mechanistic explanations for flowering schedules based on such “pollen coupling” are likely even more complex in animal-pollinated species, as they may also require accounting for the behavior and population dynamics of pollinators.

Lastly, both heritable variation in reproductive allocation and timing among individuals (Geber and Griffin, 2003; Kang et al., 2003; Santos-del-Blanco et al., 2010), as well as acquired differences that persist throughout an individual’s life (Haymes & Fox, 2012; Clark et al., 2010) are likely to be important. Dainou et al. (2012) found evidence of genetic isolation by time, but not by distance, at small spatial scales in a tropical tree—synchronously flowering individuals were more likely to be related, as well as more likely to exchange pollen. In another tropical tree, Monthe et al. found evidence of assortative mating despite extensive pollen flow, suggesting related individuals were either more likely to synchronize flowering or to be visited by the same pollinators. These intriguing results highlight how taking into account individual variation in flowering can yield new insights into the evolution and ecology of tropical trees. More broadly, long-term phenology datasets should endeavor to include information specific to individuals; this data is crucial if we are going to understand the drivers behind population- or community-level patterns in complex communities (Clark et al., 2010).

The spatial and temporal patterns of reproduction of sedentary organisms can have important implications for offspring fitness, population diversity, persistence, and species interactions in a community (Levin, 1992; Velazquez-Castro & Eichhorn, 2017). From the perspective of both population and community dynamics, flowering remains a complex and understudied aspect of the reproduction of tropical trees, yet it is a critical life history event (Abe et al., 2016.). A better understanding of both the ultimate and proximate drivers of flowering patterns will improve our understanding of biodiversity mechanisms, and is important for conservation planning in long-lived and low-density species, such as timber trees or threatened species for which genetic variation needs to be managed (Monthe et al 2017; Duminil et al. 2016). Our study highlights the great variation in flowering among and within species and suggests that there may be a relationship between reproductive status and local species abundance that does not appear to be sensitive to local biotic and abiotic environmental heterogeneity. Rarer tree species may deal with low densities by maximizing the size of their mate pool, which may help them cope with pollination despite their low numbers (Augspurger, 1981). Low proportions of flowering individuals of abundant species lead to a much narrower difference in flowering abundance among species, compared to patterns of adult relative abundance; if this is typical, it could have a stabilizing effect on species coexistence in tropical forests.

## ACKNOWLEDGEMENTS

Research and logistical support were provided by Rice University, The Smithsonian Center for Tropical Forest Science-ForestGEO, the Congo Basin Institute, and the Korup Rainforest Conservation Society; and through grants to A.P.D. by The Phipps Conservatory and ForestGEO. A.P.D. and W.M. were also supported by NSF GRFPs. We thank the Cameroon Ministry of Forests and Wildlife and the Cameroon Ministry of Scientific Research for research permission, the Missouri Botanical Garden for access to phenology records, Olivier Hardy for providing the plot phylogeny, and ForestGEO for providing access to plot census data from 2009. We are grateful to W. Asset Nkomo, R. Tchana Wandji, S. Songue, and S. Oben, for their valuable assistance with fieldwork. B. Bachelot, S. Egan, T. Miller, and three anonymous reviewers provided valuable discussion on a previous version of this manuscript.

APD conceived the ideas and designed methodology; APD collected the data; GC, DK, DWT established the KFDP; APD and MW analyzed the data; APD and AED led the writing of the manuscript. All authors contributed critically to the drafts and gave final approval for publication. All authors state they have no conflicts of interest to declare.

## Data accessibility

The data that support the findings of this study will be deposited in Dryad upon journal publication. The Korup Forest Dynamics Plot 2008-2009 ForestGeo dataset can be requested from the FDP managers: http://ctfs.si.edu/datarequest/

## Notes

http://ctfs.si.edu/datarequest/index.php/request/form/4

### Competing Interest Statement

The authors have declared no competing interest.

## REFERENCES

Abe, T., Tachiki, Y., Kon, H., Nagasaka, A., Onodera, K., Minamino, K., … Satake, A. (2016). Parameterisation and validation of a resource budget model for masting using spatiotemporal flowering data of individual trees. Ecology Letters, 19(9), 1129–1139. https://doi.org/10.1111/ele.12651

Armstrong, J. E., & Irvine, A. K. (1989). Floral Biology of Myristica insipida (Myristicaceae), a Distinctive Beetle Pollination Syndrome. American Journal of Botany, 76(1), 86. https://doi.org/10.2307/2444777

Ashman, T.-L., Knight, T. M., Steets, J. A., Amarasekare, P., Burd, M., Campbell, D. R., … others. (2004). Pollen limitation of plant reproduction: Ecological and evolutionary causes and consequences. Ecology, 85(9), 2408–2421. Retrieved from http:///doi.org/10.1890/03-8024

Asmussen, M. A. (1979). Density-Dependent Selection II. The Allee Effect. The American Naturalist, 114(6), 796–809. https://doi.org/10.1086/283529

Augspurger, C. K. (1981). Reproductive Synchrony of a Tropical Shrub: Experimental Studies on Effects of Pollinators and Seed Predators in Hybanthus Prunifolius (Violaceae). Ecology, 62(3), 775–788. https://doi.org/10.2307/1937745

Baddeley, A., Rubak, E., & Turner, R. (2015). Spatial point patterns: Methodology and applications with R. CRC Press.

Bawa, K. S., Perry, D. R., & Beach, J. H. (1985). Reproductive Biology of Tropical Lowland Rain Forest Trees. I. Sexual Systems and Incompatibility Mechanisms. American Journal of Botany, 72(3), 331. https://doi.org/10.2307/2443526

Bawa, K. S., & Webb, C. J. (1984). Flower, Fruit and Seed Abortion in Tropical Forest Trees: Implications for the Evolution of Paternal and Maternal Reproductive Patterns. American Journal of Botany, 71(5), 736–751. https://doi.org/10.2307/2443371

Bawa, Kamaljit S. (1990). Plant-pollinator interactions in tropical rain forests. Annual Review of Ecology and Systematics, 399–422. Retrieved from http://www.jstor.org/stable/2097031

Bell, J. M., Karron, J. D., & Mitchell, R. J. (2005). Interspecific Competition for Pollination Lowers Seed Production and Outcrossing in Mimulus ringens. Ecology, 86(3), 762–771. https://doi.org/10.1890/04-0694

Bella, I. E. (1971). A new competition model for individual trees. Forest Science, 17(3), 364–372. https://doi.org/10.1093/forestscience/17.3.364

Berdanier, A. B., and Clark, J. S. (2016). Divergent reproductive allocation trade-offs with canopy exposure across tree species in temperate forests. Ecosphere 7(6):e01313. http://doi.org/10.1002/ecs2.1313

Betancourt, M. (2017). A Conceptual Introduction to Hamiltonian Monte Carlo. ArXiv:1701.02434 [Stat]. Retrieved from http://arxiv.org/abs/1701.02434

Blake-Mahmud, J., & Struwe, L. (2019). Time for a change: Patterns of sex expression, health and mortality in a sex-changing tree. Annals of Botany. https://doi.org/10.1093/aob/mcz037

Bosch, M., & Waser, N. M. (1999). Effects of local density on pollination and reproduction in Delphinium nuttallianum and Aconitum columbianum (Ranunculaceae). American Journal of Botany, 86(6), 871–879. https://doi.org/10.2307/2656707

Boyle, W. A., & Bronstein, J. L. (2012). Phenology of tropical understory trees: Patterns and correlates. Revista de Biologia Tropical, 60(4), 1415–1430. https://doi.org/10.15517/RBT.V60I4.2050

Paul-Christian Bürkner (2017). brms: An R Package for Bayesian Multilevel Models Using Stan. Journal of Statistical Software, 80(1), 1–28. doi:10.18637/jss.v080.i01

Bruijning, M., Visser, M. D., Muller-Landau, H. C., Wright, S. J., Comita, L. S., Hubbell, S. P., … Jongejans, E. (2017). Surviving in a Cosexual World: A Cost-Benefit Analysis of Dioecy in Tropical Trees. The American Naturalist, 189(3), 297–314. https://doi.org/10.1086/690137

Bull, J. J., & Shine, R. (1979). Iteroparous Animals that Skip Opportunities for Reproduction. The American Naturalist, 114(2), 296–303. https://doi.org/10.1086/283476

Bush, E. R., Abernethy, K. A., Jeffery, K., Tutin, C., White, L., Dimoto, E., … Bunnefeld, N. (2016). Fourier analysis to detect phenological cycles using long-term tropical field data and simulations. Methods in Ecology and Evolution, n/a-n/a. https://doi.org/10.1111/2041-210X.12704

Canham, C. D., LePage, P. T., & Coates, K. D. (2004). A neighborhood analysis of canopy tree competition: Effects of shading versus crowding. Canadian Journal of Forest Research, 34(4), 778–787. https://doi.org/10.1139/x03-232

Carpenter, B., Gelman, A., Hoffman, M. D., Lee, D., Goodrich, B., Betancourt, M., … Riddell, A. (2017). Stan: A probabilistic programming language. Journal of Statistical Software, 76(1). https://doi.org/10.18637/jss.v076.i01

Castilla, A. R., Pope, N., & Jha, S. (2016). Positive density-dependent reproduction regulated by local kinship and size in an understorey tropical tree. Annals of Botany, 117(2), 319–329. https://doi.org/10.1093/aob/mcv170

Chesson, P. (2000). Mechanisms of Maintenance of Species Diversity. Annual Review of Ecology and Systematics, 31(1), 343–366. https://doi.org/10.1146/annurev.ecolsys.31.1.343

Chuyong, G. B., Kenfack, D., Harms, K. E., Thomas, D. W., Condit, R., & Comita, L. S. (2011). Habitat specificity and diversity of tree species in an African wet tropical forest. Plant Ecology, 212(8), 1363–1374. https://doi.org/10.1007/s11258-011-9912-4

Clark, J. S., Bell, D., Chu, C., Courbaud, B., Dietze, M., Hersh, M., … McMahon, S. (2010). High-dimensional coexistence based on individual variation: A synthesis of evidence. Ecological Monographs, 80(4), 569–608. https://doi.org/10.1890/09-1541.1

Côrtes, M. C., Uriarte, M., Lemes, M. R., Gribel, R., John Kress, W., Smouse, P. E., & Bruna, E. M. (2013). Low plant density enhances gene dispersal in the Amazonian understory herb *Heliconia acuminata*. Molecular Ecology, 22(22), 5716–5729. https://doi.org/10.1111/mec.12495

Daïnou, K., Laurenty, E., Mahy, G., Hardy, O.J., Brostaux, Y., Tagg, N. and Doucet, J.-L. (2012), Phenological patterns in a natural population of a tropical timber tree species, *Milicia excelsa* (Moraceae): Evidence of isolation by time and its interaction with feeding strategies of dispersers. American Journal of Botany, 99: 1453–1463. https://doi.org/10.3732/ajb.1200147

Dick, C. W., Hardy, O. J., Jones, F. A., & Petit, R. J. (2008). Spatial Scales of Pollen and Seed-Mediated Gene Flow in Tropical Rain Forest Trees. Tropical Plant Biology, 1(1), 20–33. https://doi.org/10.1007/s12042-007-9006-6

Dietzsch, A. C., Stanley, D. A., & Stout, J. C. (2011). Relative abundance of an invasive alien plant affects native pollination processes. Oecologia, 167(2), 469–479. https://doi.org/10.1007/s00442-011-1987-z

Duminil, J., Mendene Abessolo, D. T., Ndiade Bourobou, D., Doucet, J.-L., Loo, J., & Hardy, O. J. (2016). High selfing rate, limited pollen dispersal and inbreeding depression in the emblematic African rain forest tree Baillonella toxisperma – Management implications. Forest Ecology and Management, 379, 20–29. https://doi.org/10.1016/j.foreco.2016.08.003

Dyer, R. J., Chan, D. M., Gardiakos, V. A., & Meadows, C. A. (2012). Pollination graphs: Quantifying pollen pool covariance networks and the influence of intervening landscape on genetic connectivity in the North American understory tree, Cornus florida L. Landscape Ecology, 27(2), 239–251. https://doi.org/10.1007/s10980-011-9696-x

Egbe, E. A., Chuyong, G. B., Fonge, B. A., Namuene K. S. (2012) Forest disturbance and natural regeneration in an African raiforest at Korup National Park, Cameroon. Int. J. Biodivers. Conserv., 11, 377–384

Elzinga, J. A., Atlan, A., Biere, A., Gigord, L., Weis, A. E., & Bernasconi, G. (2007). Time after time: Flowering phenology and biotic interactions. Trends in Ecology & Evolution, 22(8), 432–439. https://doi.org/10.1016/j.tree.2007.05.006

Gabry, J., & Tristan Mahr. (2018). bayesplot: Plotting for Bayesian Models. R package version 1.5.0. Retrieved from https://CRAN.R-project.org/package=bayesplot

Geber, M. A., and Griffen, L. R., (2003). Inheritance and Natural Selection on Functional Traits. International Journal of Plant Sciences 164 (S3), S21–S42. https://doi.org/10.1086/368233

Gelman, A., Carlin, J. B., Stern, H. S., Dunson, D. B., Vehtari, A., & Rubin, D. B. (2013). Bayesian data analysis. CRC press.

Gelman, A., & Hill, J. (2007). Data analysis using regression and multilevelhierarchical models (Vol. 1). Cambridge University Press New York, NY, USA.

Ghazoul, J. (2005). Pollen and seed dispersal among dispersed plants. Biological Reviews, 80(03), 413. https://doi.org/10.1017/S1464793105006731

Grant, E.L., Conroy, G.C., Lamont, R.W. et al. Short distance pollen dispersal and low genetic diversity in a subcanopy tropical rainforest tree, *Fontainea picrosperma* (Euphorbiaceae). Heredity 123, 503–516 (2019). https://doi-org.ezproxy.rice.edu/10.1038/s41437-019-0231-1

Hardy, O. J., Maggia, L., Bandou, E., Breyne, P., Caron, H., Chevallier, M.-H., … Degen, B. (2006). Fine-scale genetic structure and gene dispersal inferences in 10 Neotropical tree species. Molecular Ecology, 15(2), 559–571. https://doi.org/10.1111/j.1365-294X.2005.02785.x

Hart, S. P., Schreiber, S. J., & Levine, J. M. (2016). How variation between individuals affects species coexistence. Ecology Letters, 19(8), 825–838. https://doi.org/10.1111/ele.12618

Harvey, P. H., & Pagel, M. D. (1991). The comparative method in evolutionary biology (Vol. 239). Oxford university press Oxford.

Haymes, K. L., & Fox, G. A. (2012). Variation among individuals in cone production in Pinus palustris (Pinaceae). American Journal of Botany, 99(4), 640–645. https://doi.org/10.3732/ajb.1100339

Hendry, A. P., & Day, T. (2005). Population structure attributable to reproductive time: Isolation by time and adaptation by time. Molecular Ecology, 14(4), 901–916. https://doi.org/10.1111/j.1365-294X.2005.02480.x

Hoch, G., Siegwolf, R. T. W., Keel, S. G., Körner, C., & Han, Q. (2013). Fruit production in three masting tree species does not rely on stored carbon reserves. Oecologia, 171(3), 653–662. https://doi.org/10.1007/s00442-012-2579-2

Hoffman, M. D., & Gelman, A. (2014). The No-U-turn sampler: Adaptively setting path lengths in Hamiltonian Monte Carlo. Journal of Machine Learning Research, 15(1), 1593–1623.

Isagi, Y., Sugimura, K., Sumida, A., & Ito, H. (1997). How Does Masting Happen and Synchronize? Journal of Theoretical Biology, 187(2), 231–239. https://doi.org/10.1006/jtbi.1997.0442

Jones F.A, & Comita L.S. (2008). Neighbourhood density and genetic relatedness interact to determine fruit set and abortion rates in a continuous tropical tree population. Proceedings of the Royal Society B: Biological Sciences, 275(1652), 2759–2767. https://doi.org/10.1098/rspb.2008.0894

Kainer, K. A., Wadt, L. H. O., & Staudhammer, C. L. (2007). Explaining variation in Brazil nut fruit production. Forest Ecology and Management, 250(3), 244–255. https://doi.org/10.1016/j.foreco.2007.05.024

Kang, K.-S., Bila, A. D., Harju, A. M., & Lindgren, D. (2003). Estimation of fertility variation in forest tree populations. Forestry: An International Journal of Forest Research, 76(3), 329–344. https://doi.org/10.1093/forestry/76.3.329

Kenfack, D., Thomas, D. W., Chuyong, G., & Condit, R. (2007). Rarity and abundance in a diverse African forest. Biodiversity and Conservation, 16(7), 2045–2074. https://doi.org/10.1007/s10531-006-9065-2

Kenta, T., Isagi, Y., Nakagawa, M., Yamashita, M., & Nakashizuka, T. (2004). Variation in pollen dispersal between years with different pollination conditions in a tropical emergent tree. Molecular Ecology, 13(11), 3575–3584. https://doi.org/10.1111/j.1365-294X.2004.02345.x

Kunin, W. E. (1997a). Population size and density effects in pollination: Pollinator foraging and plant reproductive success in experimental arrays of Brassica kaber. Journal of Ecology, 225–234. Retrieved from http://www.jstor.org/stable/2960653

Kunin, W. E. (1997b). Population Size and Density Effects in Pollination: Pollinator Foraging and Plant Reproductive Success in Experimental Arrays of Brassica Kaber. Journal of Ecology, 85(2), 225–234. https://doi.org/10.2307/2960653

Kunin, W., & Iwasa, Y. (1996). Pollinator Foraging Strategies in Mixed Floral Arrays: Density Effects and Floral Constancy. Theoretical Population Biology, 49(2), 232–263. https://doi.org/10.1006/tpbi.1996.0013

Lasky, J. R., Bachelot, B., Muscarella, R., Schwartz, N., Forero-Montaña, J., Nytch, C. J., … Uriarte, M. (2015). Ontogenetic shifts in trait-mediated mechanisms of plant community assembly. Ecology, 96(8), 2157–2169. https://doi.org/10.1890/14-1809.1

Levin, D. A., & Anderson, W. W. (1970). Competition for Pollinators between Simultaneously Flowering Species. The American Naturalist, 104(939), 455–467. https://doi.org/10.1086/282680

Levin, S. A. (1992). The Problem of Pattern and Scale in Ecology: The Robert H. MacArthur Award Lecture. Ecology, 73(6), 1943–1967. https://doi.org/10.2307/1941447

Libalah, M. B.; Droissart, V.; Sonke, B.; Hardy, O. J.; Drouet, T.; Pescador, D. S.; Kefack, D.; Thomas, D. W.; Chuyong, G. B.; Couteron, P. (2017). Shift in functional traits along soil fertility gradient reflects non-random community assembly in a tropical African rainforest. Plant Ecology and Evolution, 150(3), 265–278

MacArthur, R. H. (1958). Population Ecology of Some Warblers of Northeastern Coniferous Forests. Ecology, 39(4), 599–619. https://doi.org/10.2307/1931600

Minor, D. M., & Kobe, R. K. (2019). Fruit production is influenced by tree size and size-asymmetric crowding in a wet tropical forest. Ecology and Evolution, 9(3), 1458–1472. https://doi.org/10.1002/ece3.4867

Momose, K. (2004). Plant reproductive interval and population density in aseasonal tropics. Ecological Research, 19(2), 245–253. https://doi.org/10.1111/j.1440-1703.2003.00629.x

Momose, K., Yumoto, T., Nagamitsu, T., Kato, M., Nagamasu, H., Sakai, S., … Inoue, T. (1998). Pollination biology in a lowland dipterocarp forest in Sarawak, Malaysia. I. Characteristics of the plant-pollinator community in a lowland dipterocarp forest. American Journal of Botany, 85(10), 1477–1501. https://doi.org/10.2307/2446404

Monthe, FK, Hardy, OJ, Doucet, J-L, Loo, J, Duminil, J. Extensive seed and pollen dispersal and assortative mating in the rain forest tree *Entandrophragma cylindricum* (Meliaceae) inferred from indirect and direct analyses. Mol Ecol. 2017; 26: 5279–5291 https://doi.org/10.1111/mec.14241

Murawski, D. A., Hamrick, J. L., & others. (1991). The effect of the density of flowering individuals on the mating systems of nine tropical tree species. Heredity, 67(2), 167–174. https://doi.org/10.1038/hdy.1991.76

Nason, J., Herre, E. & Hamrick, J. (1998). The breeding structure of a tropical keystone plant resource. Nature, 391, 685–687. https://doi.org/10.1038/35607

Neal, R. M. (2011). MCMC using Hamiltonian dynamics. Handbook of Markov Chain Monte Carlo, 2(11). CRC Press.

Newstrom, L. E., Frankie, G. W., & Baker, H. G. (1994). A New Classification for Plant Phenology Based on Flowering Patterns in Lowland Tropical Rain Forest Trees at La Selva, Costa Rica. Biotropica, 26(2), 141–159. https://doi.org/10.2307/2388804

Nottebrock, H., Schmid, B., Treurnicht, M., Pagel, J., Esler, K. J., Böhning-Gaese, K., … Schurr, F. M. (2017). Coexistence of plant species in a biodiversity hotspot is stabilized by competition but not by seed predation. Oikos, 126(2), n/a-n/a. https://doi.org/10.1111/oik.03438

Obeso, J.R. (2002), The costs of reproduction in plants. New Phytologist, 155: 321–348. doi:10.1046/j.1469-8137.2002.00477.x

Osada, N., Sugiura, S., Kawamura, K., Cho, M., & Takeda, H. (2003). Community-level flowering phenology and fruit set: Comparative study of 25 woody species in a secondary forest in Japan. Ecological Research, 18(6), 711–723. https://doi.org/10.1111/j.1440-1703.2003.00590.x

Osunkoya, O. O., Omar-Ali, K., Amit, N., Dayan, J., Daud, D. S., & Sheng, T. K. (2007). Comparative height–crown allometry and mechanical design in 22 tree species of Kuala Belalong rainforest, Brunei, Borneo. American Journal of Botany, 94(12), 1951–1962. https://doi.org/10.3732/ajb.94.12.1951

Ottewell, K., Grey, E., Castillo, F., & Karubian, J. (2012). The pollen dispersal kernel and mating system of an insect-pollinated tropical palm, *Oenocarpus bataua*. Heredity, 109(6), 332–339. https://doi.org/10.1038/hdy.2012.40

Ouédraogo, D.-Y., Doucet, J.-L., Daïnou, K., Baya, F., Biwolé, A.B., Bourland, N., Fétéké, F., Gillet, J.-F., Kouadio, Y.L. and Fayolle, A. (2018), The size at reproduction of canopy tree species in central Africa. Biotropica, 50: 465–476. doi:10.1111/btp.12531

Parmentier, I., Duminil, J., Kuzmina, M., Philippe, M., Thomas, D. W., Kenfack, D., … Hardy, O. J. (2013). How Effective Are DNA Barcodes in the Identification of African Rainforest Trees? PLoS ONE, 8(4), e54921. https://doi.org/10.1371/journal.pone.0054921

Pau, S, Okamoto, DK, Calderón, O, Wright, SJ. Long-term increases in tropical flowering activity across growth forms in response to rising CO2 and climate change. Glob. Change Biol. 2018; 24: 2105–2116. https://doi-org.ezproxy.rice.edu/10.1111/gcb.14004

Pearse, I. S., Koenig, W. D., Funk, K. A., & Pesendorfer, M. B. (2015). Pollen limitation and flower abortion in a wind-pollinated, masting tree. Ecology, 96(2), 587–593. https://doi.org/10.1890/14-0297.1

Pearse, I. S., Koenig, W. D., & Kelly, D. (2016). Mechanisms of mast seeding: Resources, weather, cues, and selection. New Phytologist, 212(3), 546–562. https://doi.org/10.1111/nph.14114

Queenborough, S. A., Burslem, D. F. R. P., Garwood, N. C., & Valencia, R. (2007). Determinants of biased sex ratios and inter-sex costs of reproduction in dioecious tropical forest trees. American Journal of Botany, 94(1), 67–78. https://doi.org/10.3732/ajb.94.1.67

R Project Core Development Team. (2017). R: A language and environment for statistical computing (Version Version 3.4.3). Retrieved from http://www.R-project.org

Revell, L. J. (2012). phytools: An R package for phylogenetic comparative biology (and other things). Methods in Ecology and Evolution, 3(2), 217–223. https://doi.org/10.1111/j.2041-210X.2011.00169.x

Robin, X., Turck, N., Hainard, A., Tiberti, N., Lisacek, F., Sanchez, J.-C., & Müller, M. (2011). pROC: An open-source package for R and S+ to analyze and compare ROC curves. BMC Bioinformatics, 12(1), 77. https://doi.org/10.1186/1471-2105-12-77

Robledo-Arnuncio, J. J., & Austerlitz, F. (2006). Pollen Dispersal in Spatially Aggregated Populations. The American Naturalist, 168(4), 500–511. https://doi.org/10.1086/507881

Rosenheim, J. A., Williams, Neal M., & Schreiber, S. J. (2014). Parental Optimism versus Parental Pessimism in Plants: How Common Should We Expect Pollen Limitation to Be? The American Naturalist, 184(1), 75–90. https://doi.org/10.1086/676503

Sakai, S., Momose, K., Yumoto, T., Nagamitsu, T., Nagamasu, H., Hamid, A. A., & Nakashizuka, T. (1999). Plant reproductive phenology over four years including an episode of general flowering in a lowland dipterocarp forest,Sarawak, Malaysia. American Journal of Botany, 86(10), 1414–1436. https://doi.org/10.2307/2656924

Santos-del-Blanco, L., Zas, R., Notivol, E., Chambel, M. R., Majada, J., & Climent, J. (2010). Variation of early reproductive allocation in multi-site genetic trials of Maritime pine and Aleppo pine. Forest Systems, 19(3), 381–392. https://doi.org/10.5424/fs/2010193-9109

Sargent, R. D., & Vamosi, J. C. (2008). The Influence of Canopy Position, Pollinator Syndrome, and Region on Evolutionary Transitions in Pollinator Guild Size. International Journal of Plant Sciences, 169(1), 39–47. https://doi.org/10.1086/523359

Schauber, E.M., Kelly, D., Turchin, P., Simon, C., Lee, W.G., Allen, R.B., Payton, I.J., Wilson, P.R., Cowan, P.E. and Brockie, R.E. (2002), Masting by eighteen New Zealand plant species: the role of temperature as a synchronizing cue. Ecology, 83: 1214–1225. doi:10.1890/0012-9658(2002)083[1214:MBENZP]2.0.CO;2

Schmid, B., Nottebrock, H., Esler, K. J., Pagel, J., Pauw, A., Böhning-Gaese, K., … Schleuning, M. (n.d.). Responses of nectar-feeding birds to floral resources at multiple spatial scales. Ecography, 39(7), 619–629. https://doi.org/10.1111/ecog.01621

Sedgley, M., & Griffin, A. R. (2013). Sexual Reproduction of Tree Crops. Academic Press.

Stacy, E. A., Hamrick, J. L., Nason, J. D., Hubbell, S. P., Foster, R. B., & Condit, R. (1996). Pollen dispersal in low-density populations of three neotropical tree species. American Naturalist, 275–298. https://doi.org/10.1086/285925

Taylor, B. N., Chazdon, R. L., Bachelot, B., & Menge, D. N. L. (2017). Nitrogen-fixing trees inhibit growth of regenerating Costa Rican rainforests. Proceedings of the National Academy of Sciences, 114(33), 8817–8822. https://doi.org/10.1073/pnas.1707094114

Thomas, D. W., Kenfack, D., Chuyong, G. B., Moses, S. N., Losos, E. C., Condit, R. S., … others. (2003). Tree species of Southwestern Cameroon: Tree distribution maps, diameter tables, and species documentation of the 50-hectare Korup Forest Dynamics Plot. Center for Tropical Forest Science, the Smithsonian Tropical Forest Research Institute.

Thomas, S. C., & LaFrankie, J. V. (1993). Sex, Size and Interyear Variation in Flowering Among Dioecious Trees of the Malayan Rain Forest. Ecology, 74(5), 1529–1537. https://doi.org/10.2307/1940080

Thomas, Sean C. (1996). Relative Size at Onset of Maturity in Rain Forest Trees: A Comparative Analysis of 37 Malaysian Species. Oikos, 76(1), 145–154. https://doi.org/10.2307/3545756

Uriarte, M., Condit, R., Canham, C. D., & Hubbell, S. P. (2004). A spatially explicit model of sapling growth in a tropical forest: Does the identity of neighbours matter? Journal of Ecology, 92(2), 348–360. https://doi.org/10.1111/j.0022-0477.2004.00867.x

Vaast, P., Bertrand, B., Perriot, J.-J., Guyot, B., & Génard, M. (2006). Fruit thinning and shade improve bean characteristics and beverage quality of coffee (Coffea arabica L.) under optimal conditions. Journal of the Science of Food and Agriculture, 86(2), 197–204. https://doi.org/10.1002/jsfa.2338

Vamosi, J. C., Steets, J. A., & Ashman, T.-L. (2013). Drivers of pollen limitation: Macroecological interactions between breeding system, rarity, and diversity. Plant Ecology & Diversity, 6(2), 171–180. https://doi.org/10.1080/17550874.2013.769130

Velazquez-Castro, J., & Eichhorn, M. P. (2017). Relative ranges of mating and dispersal modulate Allee thresholds in sessile species. Ecological Modelling, 359, 269–275. https://doi.org/10.1016/j.ecolmodel.2017.05.025

Visser, M.D., Bruijning, M., Wright, S.J., Muller-Landau, H.C., Jongejans, E., Comita, L.S. and de, Kroon, H. (2016), Functional traits as predictors of vital rates across the life cycle of tropical trees. Funct Ecol, 30: 168–180. https://doi.org/10.1111/1365-2435.12621

Waples, R. S., & Antao, T. (2014). Intermittent Breeding and Constraints on Litter Size: Consequences for Effective Population Size per Generation (Ne) and per Reproductive Cycle (Nb). Evolution, 68(6), 1722–1734. https://doi.org/10.1111/evo.12384

Whitney, K. D. (2009). Comparative Evolution of Flower and Fruit Morphology. Proceedings: Biological Sciences, 276(1669), 2941–2947. https://doi.org/10.1098/rspb.2009.0483

Wickham, H. (2016). ggplot2: Elegant Graphics for Data Analysis. Springer.

Winn, A., Elle, E., Kalisz, S., Cheptou, P.-O., Eckert, C. G., Goodwillie, C., … Vallejo-Marín, M. (2011). Analysis of Inbreeding Depression in Mixed-Mating Plants Provides Evidence for Selective Interference and Stable Mixed Mating. Evolution, 65(12), 3339–3359. https://doi.org/10.1111/j.1558-5646.2011.01462.x

Wolkovich, E. M., Cook, B. I., McLauchlan, K. K., & Davies, T. J. (2014). Temporal ecology in the Anthropocene. Ecology Letters, 17(11), 1365–1379. https://doi.org/10.1111/ele.12353

Wright, S. J., Jaramillo, M. A., Pavon, J., Condit, R., Hubbell, S. P., & Foster, R. B. (2005). Reproductive size thresholds in tropical trees: Variation among individuals, species and forests. Journal of Tropical Ecology, 21(3), 307–315. https://doi.org/10.1017/S0266467405002294

