## Supplementary material for "Ecological correlates of reproductive status in a guild of Afrotropical understory trees": Fig. S1

### SUPPORTING INFORMATION

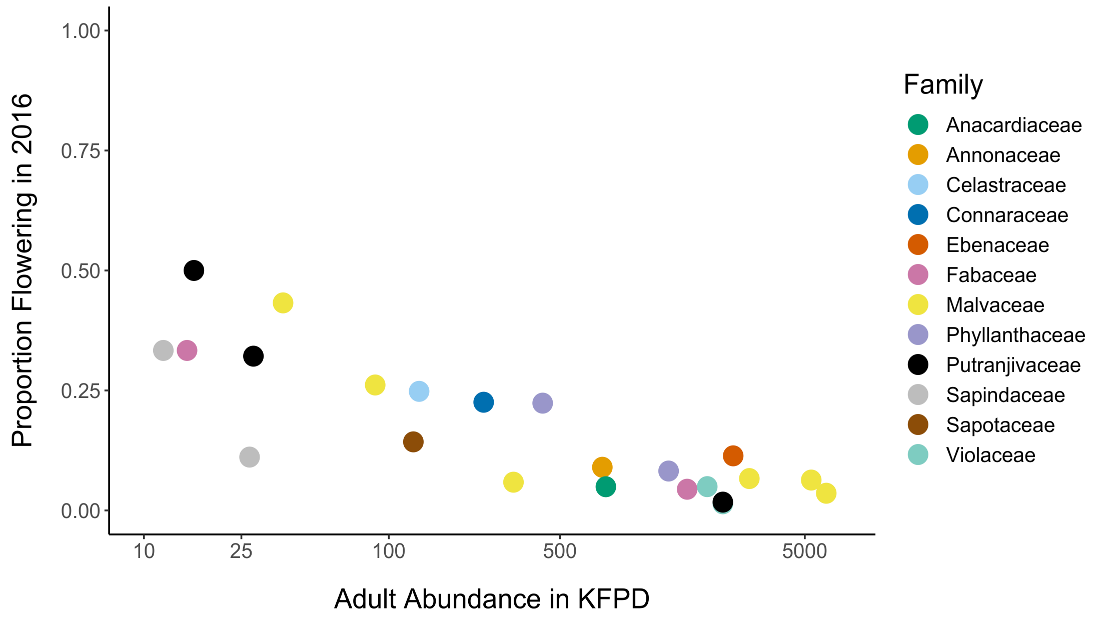


**Figure S1.** Proportion of flowering individuals compared to adult abundance in 25ha. Dots represent species, colors represent taxonomic families. Note negative trend with increasing adult abundance.
