## Supplementary material for "Ecological correlates of reproductive status in a guild of Afrotropical understory trees": Fig. S2

### SUPPORTING INFORMATION

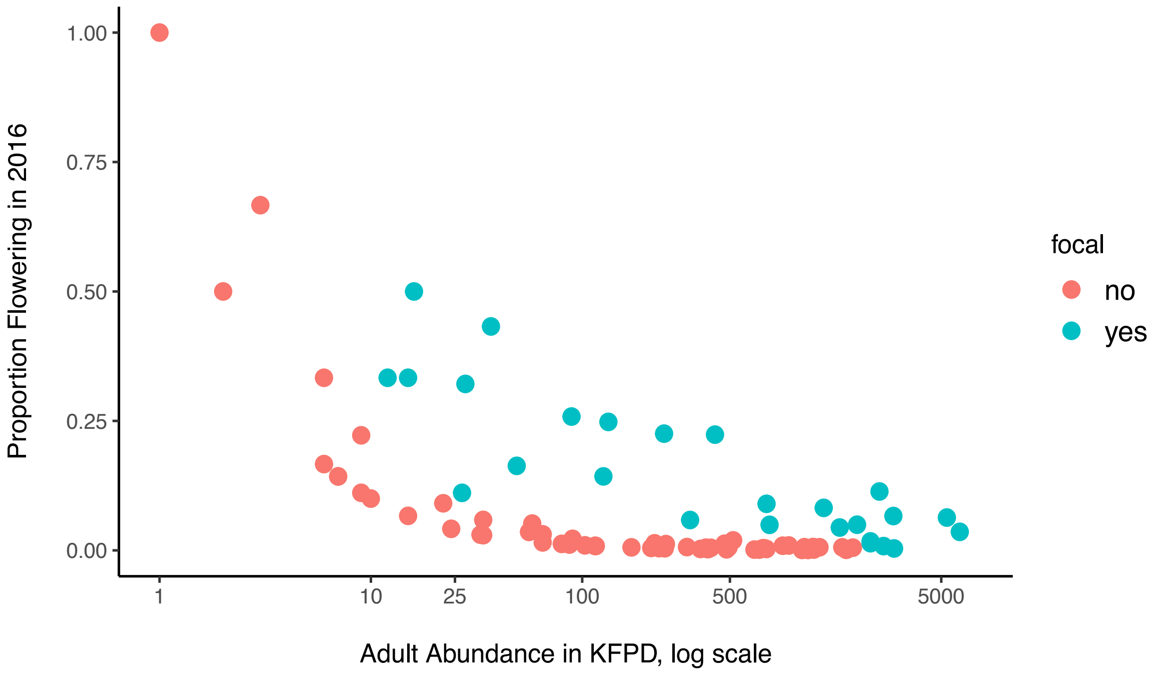


**Figure S2.** Proportion of flowering individuals of focal species and non-focals (those that were either too rare, had < 3 individuals flowering or may have been under-sampled for other reasons) compared to adult abundance, log scale, in 25ha.
