## Supplementary material for "Ecological correlates of reproductive status in a guild of Afrotropical understory trees": Fig. S3

### SUPPORTING INFORMATION

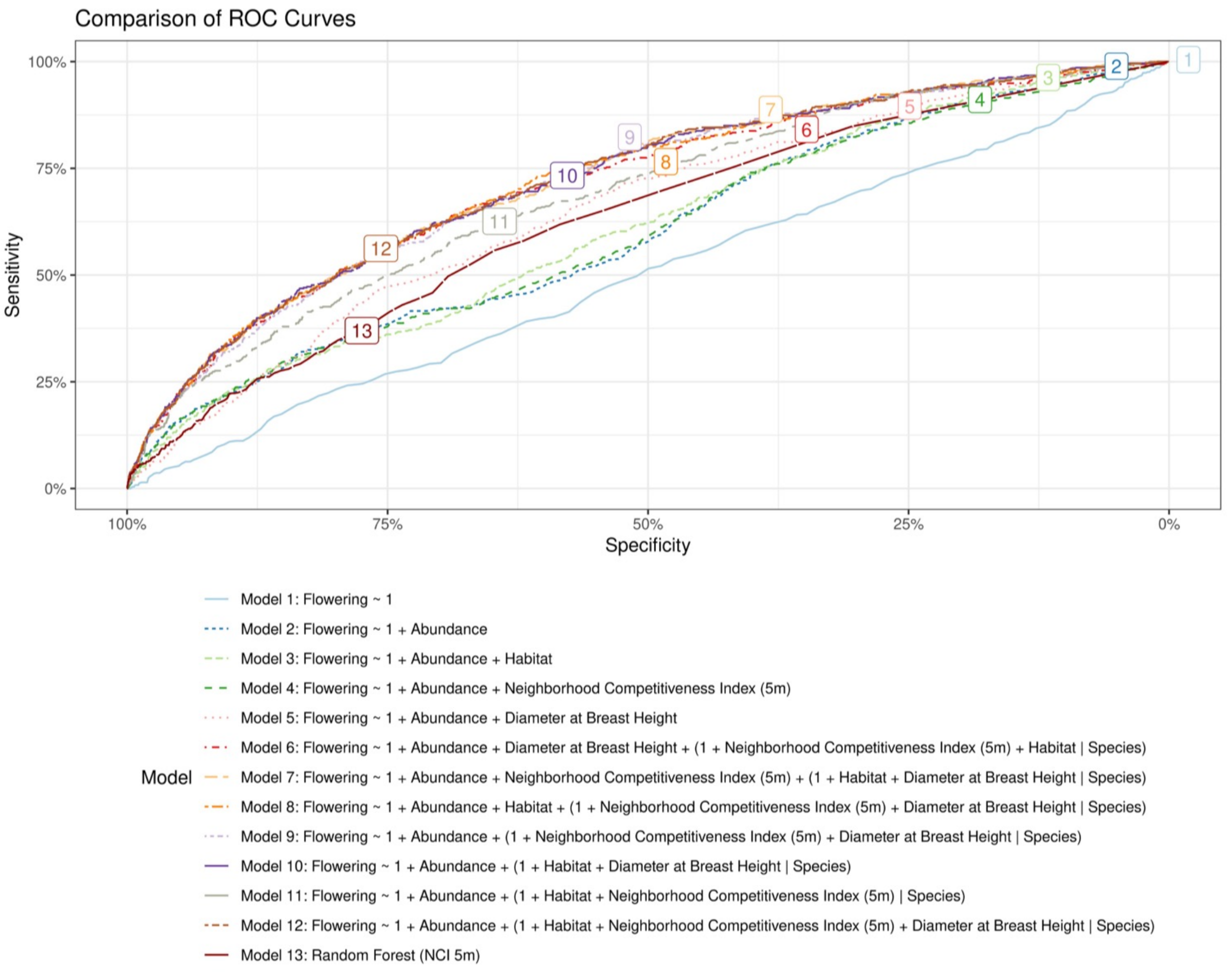


**Figure S3.** Comparison of the (out of sample) ROC curves for many of the models evaluated. The best performing models all outperform a random forest model and have per-species random intercepts and random slopes for DBH and NCI.
