## Supplementary material for "Ecological correlates of reproductive status in a guild of Afrotropical understory trees": Fig. S4

### SUPPORTING INFORMATION

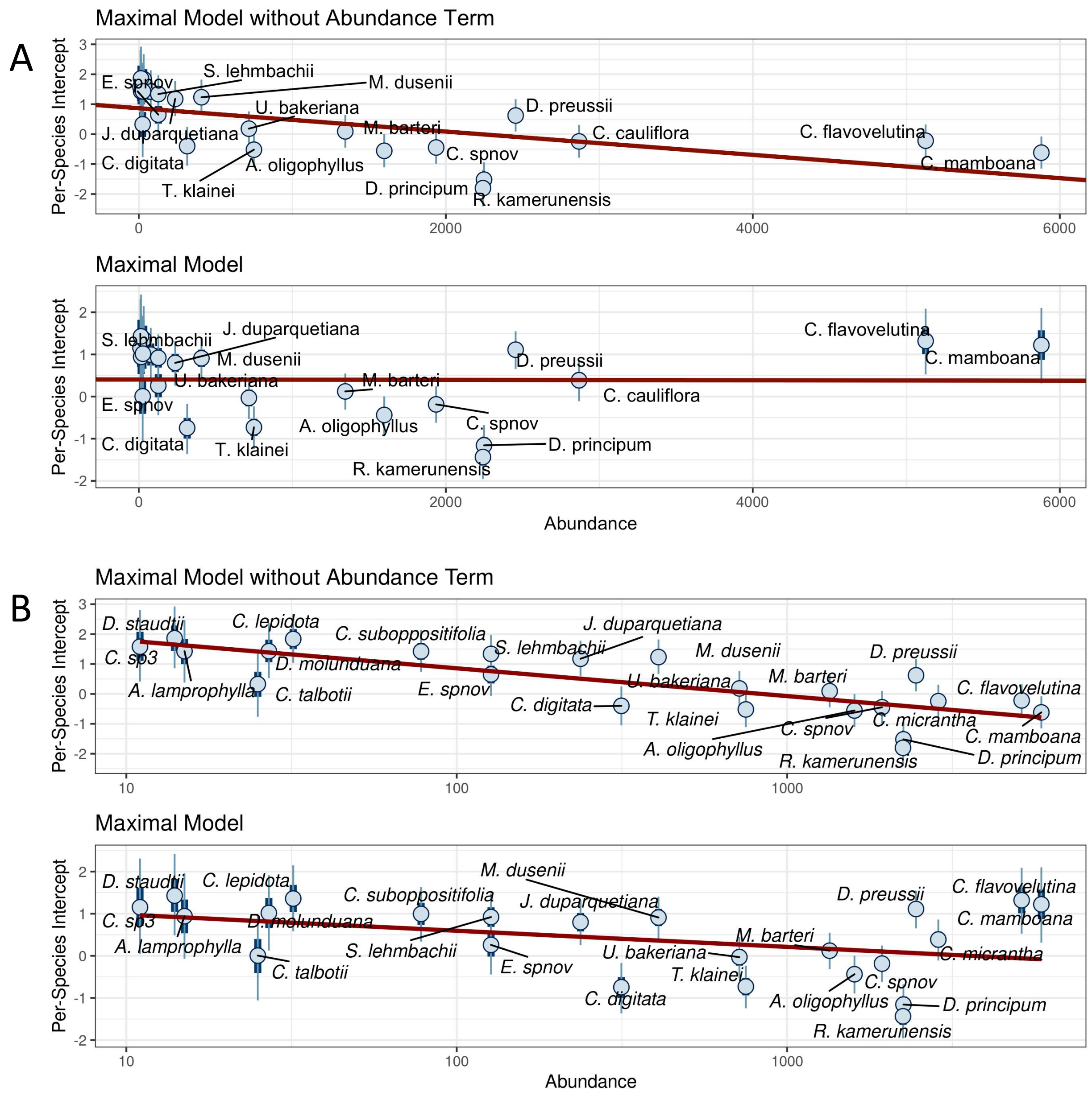


**Figure S4.** Species-specific intercepts from the maximal model with and without the abundance covariate included. The red line is the least squares (OLS) line of the estimated per species effect plotted against per-species abundances. When the abundance covariate is omitted (top), there is a strong negative correlation between the estimated per-species intercepts and abundance. By contrast, when the abundance covariate is included (below), the correlation between intercept and abundance disappears and the standard deviation of the per-species intercepts is decreased. This suggests that abundance could be a driver of the observed per-species variation, even after controlling for habitat, size, and local crowding. The bottom panel is the best-performing model (Model 18) while the top panel is Model 18 with the explicit abundance effect removed. Top two plots shows Abundance, while bottom two plots show log-transformed Abundance on the x-axis.
