## Supplementary material for "Ecological correlates of reproductive status in a guild of Afrotropical understory trees": Fig. S5

### SUPPORTING INFORMATION

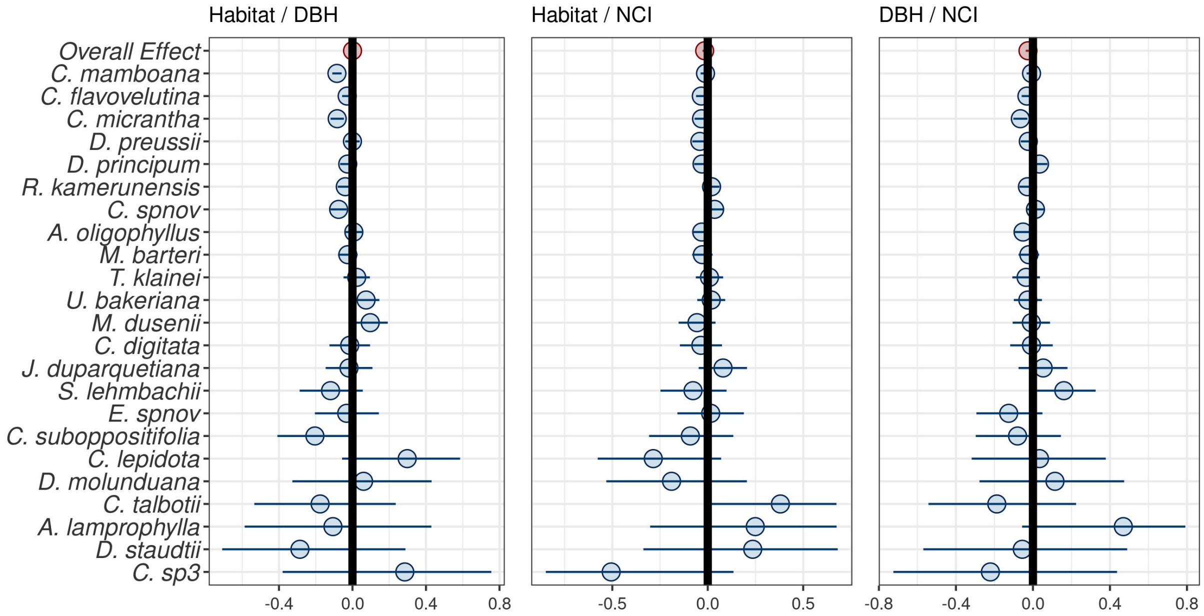


**Figure S5.** Pearson Correlations between Diameter at Breast Height (DBH) and Neighborhood Competitive Index (NCI) (5m radius) (left panel), Habitat and NCI (5m radius) (middle panel), DBH and NCI (right panel).
