## Supplementary material for "Ecological correlates of reproductive status in a guild of Afrotropical understory trees": Fig. S6

**SUPPORTING INFORMATION**

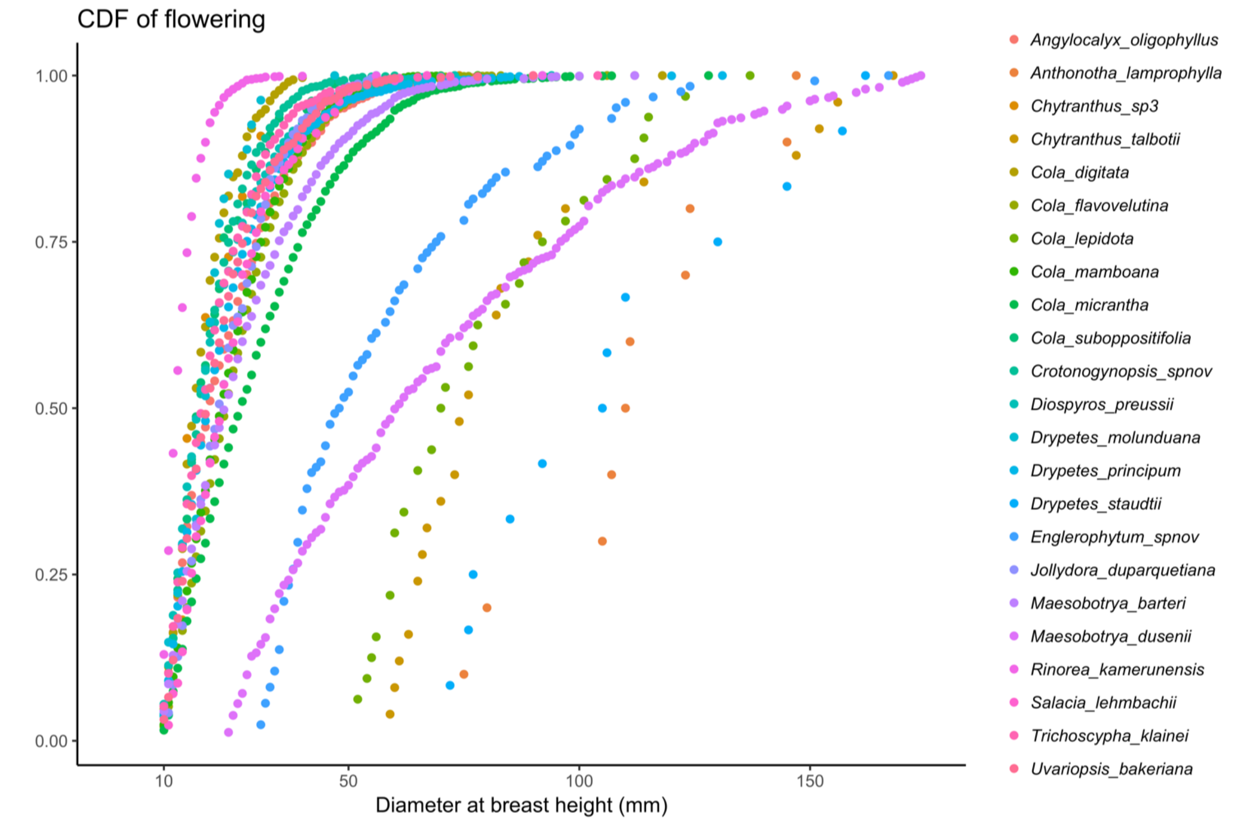


**Figure S6.** Cumulative distribution plot (CDF) of diameter at breast height (DBH) for flowering focal species, 2016. Note how larger sized species have wider variation in size at reproduction.
