## Supplementary material for "Ecological correlates of reproductive status in a guild of Afrotropical understory trees": Table S1

### SUPPORTING INFORMATION

**Table S1.** Focal species information

| Family | Species | KFDP ForestGEO code | Life Form (T=treelet; ST=small tree; MT: medium tree) | N total adults 2009 census | N reproductives 2016 | Minimum reproductive** DBH (mm) | Reproductive nearest consp. dist. significantly overdispersed relative to adult random sample NCD? | Reproductive Morphology | Relative fruit size/seed size (from Thomas et al. 2003) | Fleshy/not | Capable of self-fertilization (with or without pollinator assistance) |
|---|---|---|---|---|---|---|---|---|---|---|---|
| Anacadriaceae | *Trichoschypha klainei* | TRI3 | ST | 770 | 38 | 10 | yes | dioecious | medium | Fleshy | no |
| Annonaceae | *Uvariopsis bakeriana* | UVBA | T | 745 | 67 | 10 | yes | polygamodioecious | large/small | Fleshy | ? |
| Celastraceae | *Salacia lehmbachii* | SAL2 | T | 133 | 33 | 11 | no | bisexual, protandrous | large/large | Fleshy | ? |
| Connaraceae | *Jollydora duparquetiana* | JOLY | ST | 244 | 55 | 11 | yes | andromonoecious | large/large | Fleshy | yes?* |
| Ebenaceae | *Diospyros preussii* | DIPR | ST | 2550 | 290 | 10 | yes | polygamodioecious | medium/medium | Fleshy | ? |
| Euphorbiaceae | *Crotonogynopsis korupensis* | ALEX | T | 1998 | 99 | 10 | yes | dioecious | ? | Not | no |
| Fabaceae | *Anthonotha lamprophylla* | ANLA | ST | 15 | 5 | 74 | no | bisexual | ? | Not? | ? |
| Fabaceae | *Angylocalyx oligophyllus* | ANTA | T | 1654 | 73 | 10 | yes | bisexual | medium/medium | Fleshy | ? |
| Malvaceae | *Cola lepidota* | COLE | ST | 37 | 16 | 52 | yes | monoecious | large/large | Fleshy | ? |
| Malvaceae | *Cola suboppositofolia* | CONS | T | 88 | 23 | 11 | no | monoecious | large/large | Fleshy | ? |
| Malvaceae | *Cola digitata* | CODI | T | 323 | 19 | 10 | no | monoecious | large/large | Fleshy | ? |
| Malvaceae | *Cola micrantha* | COCA | ST | 2968 | 197 | 10 | yes | monoecious | large/large | Fleshy | ? |
| Malvaceae | *Cola flavovelutina* | OCTI | ST | 5312 | 336 | 10 | yes | monoecious | large/large | Fleshy | ? |
| Malvaceae | *Cola mamboana* | COAT | T | 6118 | 219 | 10 | yes | monoecious | large/large | Fleshy | no |
| Phyllanthaceae | *Maesobotrya dunsenii* | MADU | ST | 425 | 95 | 24 | yes | dioecious | medium/tiny | Fleshy | no |
| Phyllanthaceae | *Maesobotrya barterii* | MABA | T | 1389 | 114 | 10 | yes | dioecious | small/tiny | Fleshy | no |
| Putranjivaceae | *Drypetes molonduana* | DRY3 | T | 28 | 9 | 10 | no | dioecious | ? | Fleshy | no |
| Putranjivaceae | *Drypetes staudtii* | DRST | MT | 16 | 8 | 71 | no | dioecious | large/ | Fleshy | no |
| Putranjivaceae | *Drypetes principum* | DRSI2 | ST | 2313 | 40 | 10 | yes | dioecious | large/ | Fleshy | no |
| Sapindaceae | *Chytranthus sp.3* | CHY4 | T | 12 | 4 | 13 | no | andromonoecious | ? | Fleshy | ? |
| Sapindaceae | *Chytranthus talbotii* | CHTA | ST | 27 | 3 | 59 | no | probably dioecious? | ? | Fleshy | ? |
| Sapotaceae | *Englerophytum sp.nov.* | ENGI | ST | 126 | 18 | 31 | yes | bisexual | large/? | Fleshy | ? |
| Violaceae | *Rinorea kamerunensis* | RINA | T | 2313 | 32 | 10 | yes | bisexual | small/small | Not | ? |

* We performed an unassisted selfing experiment (unpublished) by excluding all floral visitors on one individual of *Jollydora duparquetiana* and found positive seed set similar to open-pollinated individuals.

** Based on smallest flowering DBH observed. See also Figure S5 for cumulative density curves of flowering DBH for 2016
