## Supplementary material for "Ecological correlates of reproductive status in a guild of Afrotropical understory trees": Table S2

### SUPPORTING INFORMATION

**Table S2.** Models and comparison of predictive power using area under the curve/ROC analysis. Model 18 (in bold) was selected as the best model, as it has the best out-of-sample predictive performance and treats all predictors (NCI, DBH, and Habitat) symmetrically. Models are specified using the right hand side (predictors) of the formula notation of R's lme4 package for mixed-effects models.

| Model | Model Specification | Mean AUC | Standard Error |
|---|---|---|---|
| 1 | 1 | 0.511 | 0.004 |
| 2 | 1 + Abundance | 0.578 | 0.006 |
| 3 | 1 + Abundance + Habitat | 0.581 | 0.008 |
| 4 | 1 + Abundance + Neighborhood Competitiveness Index (5m) | 0.582 | 0.008 |
| 5 | 1 + Abundance + Diameter at Breast Height | 0.634 | 0.004 |
| 6 | 1 + Abundance + Neighborhood Competitiveness Index (5m) + Diameter at Breast [illegible] | 0.634 | 0.004 |
| 7 | 1 + Abundance + Neighborhood Competitiveness Index (5m) + Habitat | 0.582 | 0.007 |
| 8 | 1 + Abundance + Habitat + Diameter at Breast Height | 0.633 | 0.004 |
| 9 | 1 + Abundance + Neighborhood Competitiveness Index (5m) + Diameter at Breast Height + Habitat | 0.632 | 0.004 |
| 10 | 1 + Abundance + Neighborhood Competitiveness Index (5m) + Diameter at Breast Height + (1 + Habitat \| Species) | 0.705 | 0.005 |
| 11 | 1 + Abundance + Habitat + Neighborhood Competitiveness Index (5m) + Diameter at Breast Height + (1 \| Species) | 0.704 | 0.005 |
| 12 | 1 + Abundance + Habitat + Neighborhood Competitiveness Index (5m) + Diameter at Breast Height + (1 \| Phylogeny) | 0.703 | 0.005 |
| 13 | 1 + Abundance + Neighborhood Competitiveness Index (5m) + Diameter at Breast Height + (1 + Habitat \| Species) | 0.704 | 0.005 |
| 14 | 1 + Abundance + Habitat + Diameter at Breast Height + (1 + Neighborhood Competitiveness Index (5m) \| Species) | 0.704 | 0.005 |
| 15 | 1 + Abundance + Diameter at Breast Height + (1 + Neighborhood Competitiveness Index (5m) + Habitat \| Species) | 0.705 | 0.005 |
| 16 | 1 + Abundance + Habitat + Neighborhood Competitiveness Index (5m) + (1 + Diameter at Breast Height \| Species) | 0.712 | 0.005 |
| 17 | 1 + Abundance + Neighborhood Competitiveness Index (5m) + (1 + Habitat + Diameter at Breast Height \| Species) | 0.713 | 0.005 |
| 18 | 1 + Abundance + (1 + Habitat + Neighborhood Competitiveness Index (5m) + Diameter at Breast Height \| Species) | 0.713 | 0.005 |
| 19 | 1 + Abundance + (1 + Habitat + Neighborhood Competitiveness Index (5m) + Diameter at Breast Height \| Phylogeny) | 0.711 | 0.006 |
| 20 | 1 + Abundance + Habitat + (1 + Neighborhood Competitiveness Index (5m) + Diameter at Breast Height \| Species) | 0.713 | 0.005 |
| 21 | 1 + Abundance + (1 + Habitat + Neighborhood Competitiveness Index (10m) + Diameter at Breast Height \| Species) | 0.712 | 0.005 |
| 22 | 1 + Abundance + (1 + Habitat + Neighborhood Competitiveness Index (15m) + Diameter at Breast Height \| Species) | 0.707 | 0.005 |
| 23 | 1 + Abundance + (1 + Neighborhood Competitiveness Index (5m) + Diameter at Breast Height \| Species) | 0.712 | 0.005 |
| 24 | 1 + Abundance + (1 + Habitat + Diameter at Breast Height \| Species) | 0.713 | 0.005 |
| 25 | 1 + Abundance + (1 + Habitat + Neighborhood Competitiveness Index (5m) \| Species) | 0.677 | 0.007 |
| 26 | 1 + (1 + Habitat + Neighborhood Competitiveness Index (5m) + Diameter at Breast Height \| Species) | 0.712 | 0.006 |
| 27 | Random Forest (NCI-5m) | 0.643 | 0.003 |
| 28 | Random Forest (NCI-10m) | 0.642 | 0.003 |
| 29 | Random Forest (NCI-15m) | 0.621 | 0.003 |
