## Supplementary material for "Ecological correlates of reproductive status in a guild of Afrotropical understory trees": Table S3

### SUPPORTING INFORMATION

**Table S3.** Counts for all species recorded budding and/or flowering in the understory in 2016. NB: As discussed in text, counts for non-focal species may not be accurate. PHSP and COSE were undercounted (PHSP had many <1cm (untagged) reproductive individuals and COSE peak budding phenology occurred at the end of the field study.

| KFDP code | All stems in lower 25ha 2009 ForestGEO census | Adults_2016 (based on minmimum flowering DBH 2016) | Flowering in 2016 | Focal species? | KFDP code | All stems in lower 25ha 2009 ForestGEO census | Adults_2016 (based on minmimum flowering DBH 2016) | Flowering in 2016 | Focal species? |
|---|---|---|---|---|---|---|---|---|---|
| ACI2 | 66 | 7 | 1 | no | TRAC | 15 | 6 | 2 | no |
| ACST | 42 | 34 | 1 | no | CHTA | 246 | 27 | 3 | yes |
| ANMA | 42 | 33 | 1 | no | DABL | 855 | 726 | 3 | no |
| AUTA | 1074 | 212 | 1 | no | DRLA | 252 | 220 | 3 | no |
| BEI2 | 695 | 688 | 1 | no | MIPU | 493 | 493 | 3 | no |
| BEIS | 375 | 363 | 1 | no | PANO | 58 | 58 | 3 | no |
| CHY2 | 26 | 9 | 1 | no | RUBP | 396 | 249 | 3 | no |
| CHY3 | 320 | 171 | 1 | no | SCAI | 845 | 716 | 3 | no |
| COL2 | 151 | 24 | 1 | no | CHY4 | 15 | 12 | 4 | yes |
| COLSL | 2 | 1 | 1 | no | CRAR | 1797 | 1797 | 4 | no |
| CRST | 2414 | 394 | 1 | no | ANLA | 87 | 15 | 5 | yes |
| HEPA | 653 | 653 | 1 | no | DRSI | 625 | 471 | 6 | no |
| HUUM | 58 | 2 | 1 | no | RIN6 | 1163 | 1126 | 7 | no |
| LAAF | 290 | 115 | 1 | no | DRST | 120 | 16 | 16 | yes |
| MALO | 183 | 116 | 1 | no | MAAC | 1989 | 1327 | 8 | no |
| OXY2 | 80 | 80 | 1 | no | POPA | 1270 | 1238 | 8 | no |
| PHSP2 | 1325 | 1097 | 1 | no | UOI | 1122 | 889 | 8 | no |
| PIPI | 566 | 392 | 1 | no | UVAI | 192 | 49 | 8 | yes |
| PLEI | 13 | 10 | 1 | no | BEWE | 949 | 949 | 9 | no |
| PLTA | 22 | 6 | 1 | no | DRY3 | 28 | 28 | 9 | yes |
| POMA | 410 | 231 | 1 | no | CORO | 2021 | 1909 | 10 | no |
| PSYI | 608 | 482 | 1 | no | GABI | 518 | 518 | 10 | no |
| RHA2 | 156 | 87 | 1 | no | RIN2 | 2210 | 1700 | 10 | no |
| RIN9 | 638 | 87 | 1 | no | RILE | 2990 | 2990 | 11 | yes |
| SALI | 141 | 115 | 1 | no | COLE | 76 | 37 | 16 | yes |
| SALS | 37 | 15 | 1 | no | ENGI | 196 | 126 | 18 | yes |
| TRI4 | 189 | 103 | 1 | no | CODI | 324 | 324 | 18 | yes |
| TRIC | 99 | 65 | 1 | no | RINI | 3117 | 2668 | 22 | yes |
| VOAI | 246 | 246 | 1 | no | CONS | 89 | 89 | 22 | yes |
| WAME | 1286 | 1170 | 1 | no | RINA | 2313 | 2313 | 32 | yes |
| WAR2 | 143 | 7 | 1 | no | SAL2 | 133 | 133 | 33 | yes |
| BELI | 1414 | 1247 | 2 | no | PHSP | 15734 | 15734 | 35 | no |
| CALU | 313 | 313 | 2 | no | TRI3 | 770 | 770 | 38 | yes |
| COCH | 96 | 65 | 2 | no | DRSI2 | 2313 | 2313 | 40 | yes |
| COF2 | 69 | 9 | 2 | no | JOLY | 255 | 244 | 55 | yes |
| COLS | 28 | 22 | 2 | no | UVBA | 745 | 745 | 67 | yes |
| DRYS | 105 | 56 | 2 | no | ANTA | 1654 | 1654 | 73 | yes |
| LEPA | 230 | 65 | 2 | no | MADU | 605 | 425 | 95 | yes |
| OLAI | 41 | 34 | 2 | no | ALEX | 1998 | 1998 | 99 | yes |
| OURA | 247 | 90 | 2 | no | MABA | 1389 | 1389 | 114 | yes |
| RIN3 | 1903 | 1775 | 2 | no | COSE | 11129 | 11064 | 153 | no |
| SCMA | 1017 | 744 | 2 | no | COCA | 2968 | 2968 | 197 | yes |
| TACR | 675 | 385 | 2 | no | COAT | 6118 | 6118 | 213 | yes |
| TRI2 | 586 | 406 | 2 | no | DIPR | 2550 | 2550 | 290 | yes |
| OMPI | 9 | 3 | 2 | no | OCTI | 5314 | 5314 | 337 | yes |
