## Supplementary material for "Ecological correlates of reproductive status in a guild of Afrotropical understory trees": SI7

Analysis code for section: ‘Ecological correlates of flowering status’

```
library(readr)
library(brms)
library(dplyr)
library(ape)
library(MCMCglmm)
library(foreach)
library(doMC)

registerDoMC(15)

ALL_DATA <- read_csv("all_data.csv")

phylo <- read.nexus("kphy_DRY3_APD2017.nex")

inv.phylo <- MCMCglmm::inverseA(phylo, nodes = "TIPS", scale = TRUE)
A <- solve(inv.phylo$Ainv)
colnames(A) <- rownames(A) <- rownames(inv.phylo$Ainv)

source("formulas.R", echo=TRUE)

set.seed(125)

N_SPLITS <- 5
N_FORMULAS <- length(FORMULAS)

if(!file.exists("splits.RData")){
  SPLITS <- replicate(N_SPLITS, {
    TEST_DATA <- ALL_DATA %>% group_by(code) %>% sample_frac(0.3)
    TRAIN_DATA <- setdiff(ALL_DATA, TEST_DATA)
    list(TEST=TEST_DATA, TRAIN=TRAIN_DATA)
  }, simplify=FALSE)

  saveRDS(SPLITS, "splits.RData")
} else {
  SPLITS <- readRDS("splits.RData")
}

## Fit to whole data set
foreach(f=seq_along(FORMULAS)) %dopar% {
  nm <- paste0("f_", f, "_fit.RData")
  if(!file.exists(nm)){
    cat("Starting f = ", f, "\n")
    names <- syms(intersect(unique(gsub("[()l~+-]+", "",
strsplit(paste(deparse(FORMULAS[[f]]), collapse=""), " ")[[1]])), colnames(ALL_DATA))
    this_data <- select_(ALL_DATA, .dots = names)
    this_data <- this_data[!matrixStats::rowAnys(sapply(this_data, is.infinite)),]
    if(grepl("phylo", deparse(FORMULAS[[f]]))){
```
